# Cryo-EM structures of the YsaN ATPase reveal dodecameric and tetradecameric assemblies in the *Yersinia enterocolitica* T3SS

**DOI:** 10.64898/2026.10.28.762456

**Authors:** Bhaskar Bhattacharyya, Bidisha Chakraborty, Soumyajit Nandy, Saumen Datta, Dhabaleswar Patra

## Abstract

Bacterial pathogens deploy sophisticated nanomachines such as the Type III Secretion System (T3SS) to inject effector proteins directly into host cells, thereby subverting cellular defenses and promoting infection. Central to this machinery is a conserved homo-oligomeric ATPase that fuels substrate unfolding and translocation. Despite extensive structural and biochemical studies, the precise mechanism of T3SS ATPase oligomer assembly and regulation remains incompletely understood, events crucial for the overall functionality of the T3SS. Here, we report high-resolution cryo-electron microscopy (cryo-EM) structures of the *Yersinia enterocolitica* T3SS ATPase YsaN, captured in two previously uncharacterized oligomeric states, a dodecamer (12-mer) at 3.51 Å and a tetradecamer (14-mer) at 3.80 Å resolution. Both assemblies exhibit helical-like symmetry and conformational heterogeneity. A conserved longitudinal fissure interrupts the helical lattice and appears to mediate interconversion between the two states, while an N-terminal interface stabilized by inter-tier disulfide bonds and an extensive non-covalent network locks the assembly together. Molecular dynamics simulations reveal that ATP hydrolysis drives a global, correlated compaction upon transition to the ADP-bound state. Together, these structural, biochemical, and MD simulation data establish T3SS ATPases as dynamic, oligomerically plastic molecular motors and provide a mechanistic framework for chaperone-effector dissociation and substrate translocation.

**Significance Statement:** Gram-negative bacterial pathogens utilize T3SS to actively deliver effector proteins into host cells. A conserved ATPase at its base provides the energy required for chaperone dissociation, unfolding of the effector, and driving translocation of the effector in semi-folded states. However, the mechanisms driving ATPase oligomerization remain incompletely understood. Here, we report unique 12-mer and 14-mer assemblies of YsaN, a T3SS ATPase from *Yersinia enterocolitica*. Both assemblies are stabilized by their extreme N-terminal domain, missing in all previously reported structures. Furthermore, biophysical and MD simulations revealed the dynamic nature of YsaN, providing insights into the potential mechanism underlying effector translocation. Together, these findings advance our understanding of T3SS ATPase assembly and dynamics, highlighting potential avenues for anti-virulence strategies.

## Introduction

Bacterial pathogens have evolved diverse tactics to infiltrate, inhabit, and destroy eukaryotic cells. The Type III Secretion System (T3SS) is a molecular syringe used by many pathogenic Gram-negative bacteria, including *Shigella*, *Salmonella*, enteropathogenic *Escherichia coli* (EPEC), *Yersinia*, and *Pseudomonas* species, to transfer effector proteins directly into eukaryotic host cells^1–7^. This specialized nanomachine contributes significantly to bacterial pathogenicity by altering cellular functions and impairing host defensive responses^1,4,8^. The T3SS has been established as a critical virulence determinant and, due to its non-essential role in bacterial survival, is being considered as an attractive target for anti-virulence therapy options^9–11^. From a structural perspective, the injectisome is anchored by the needle complex (NC), a multiring base that spans both bacterial membranes. This base encloses the inner membrane (IM) export apparatus^12,13^ and links to a cytoplasmic sorting platform^14^, collectively creating a continuous conduit connecting the bacterial cytosol directly to the target host cell through a translocon pore embedded in the host membrane^15–18^. Due to their shared evolutionary history, the bacterial flagellum and the T3SS conserve many fundamental structural components^19,20^.

At the centre of this secretion process lies a highly conserved cytosolic ATPase, which provides the essential mechanochemical energy required to recognize, dissociate effector-chaperone complexes, unfold substrates for transport through the narrow (∼2.5 nm diameter) channel, and energize their injection^5,21–25^. Several T3SS ATPase structures from various organisms have been reported, including InvC and SsaN of *Salmonella* ^26,27^, Spa47 of *Shigella*^28–30^, EscN of EPEC^31,32^, YscN of *Yersinia pestis*^33^, and the flagellar T3SS ATPase FliI of *Salmonella*^34,35^. These ATPases share significant sequence identity with one another^36,37^ (Fig. S4). All the reported high-resolution structures were severely limited by their missing N-terminus region. To improve solubilization, stability, and crystallizability, constructs used in all previous studies had N-terminal domains partially or completely deleted. The N-terminal domain of T3SS ATPases drives oligomerization^38^. As a result, barring the homehexameric structure of EscN (PDB ID: 6NJO), all other reported structures are monomeric, leaving a fundamental gap in our understanding of the functional oligomeric states of these ATPases.

*Yersinia enterocolitica*, a psychrophilic zoonotic pathogen, harbors two independent T3SSs: the plasmid-encoded Ysc-Yop T3SS and the chromosomally encoded Ysa-Ysp T3SS, which is primarily associated with the gastrointestinal phase of infection^39,40^. YsaN is the central ATPase of the Ysa-Ysp system.

Previous studies have reported that YsaN forms a dodecameric assembly in solution, which is unique from the standard hexameric assemblies reported for most of the T3SS ATPases^38,41^. Biochemical evidence also suggested that YsaN achieves its highest catalytic activity when assembled into higher-order oligomers (presumably dodecameric)^41^, which have previously been hypothesized to form via the stacking of two planar hexameric rings ^23,38,41,42^. Mechanistic validation of this assumption requires the full-length protein structure, the absence of which leaves a critical gap in our knowledge of the T3SS firing mechanism.

Here, we report the first full-length cryo-electron microscopy (cryo-EM) structures of any T3SS ATPase. Our analysis of intact YsaN reveals two distinct, highly active oligomeric assemblies: a dodecameric (12-mer) and a tetradecameric (14-mer) form, resolved at 3.51 Å and 3.80 Å, respectively. Crucially, these higher-order states do not form planar stacked rings, but rather assemble into dynamic, asymmetric helical-like rings containing a distinct longitudinal fissure. Supported by comprehensive ATPase assays and isothermal titration calorimetry (ITC), we demonstrate that these helical architectures represent the catalytically active states. Furthermore, using molecular dynamics (MD) simulations, we map the mechanochemical cycle of YsaN, revealing that the transition from an open, ATP-bound state to an ADP-bound state triggers structural compaction. Together, our cryo-EM, biochemical, and MD simulation data redefine T3SS ATPases as processive unfoldases, establishing the mechanical framework required for chaperone-effector dissociation, substrate unfolding, and active transport.

## Results

### Overall Structure of the Dodecameric and Tetradecameric YsaN Assemblies

Previous structural studies of T3SS ATPases have predominantly revealed canonical hexameric organizations^31^, despite biochemical evidence suggesting that YsaN natively forms higher-order oligomers^38,41^. To experimentally evaluate its assembly state, we first examined purified, full-length YsaN using negative-stain electron microscopy. This revealed double-tiered particles indicative of higher-order architectures (Fig. S1), prompting high-resolution structural characterization via single-particle cryo-EM.

Heterogeneous refinement revealed two predominant oligomeric species corresponding to a dodecameric (12-mer) and a tetradecameric (14-mer) assembly. Refinement without imposed symmetry (C1) yielded a 3.51 Å resolution reconstruction for the 12-mer and a 3.80 Å resolution reconstruction for the 14-mer (Fig. 1, Fig. S2,3, Movie S1, Table 1). The dodecameric assembly adopts a cylindrical structure comprising six dimers in which the two monomers are stacked on top of each other by their N-terminal regions. This continuous helical, double-layered cylindrical architecture has an overall diameter of ∼119 Å and a length of ∼224 Å. Similarly, the cylindrical tetradecamer comprises two helical tiers of seven protomers each, measuring ∼125 Å in diameter and ∼224 Å in length. Notably, outside of the hexameric EscN, this represents the first structural observation of a higher-order T3SS ATPase oligomer.

**Fig. 1.**
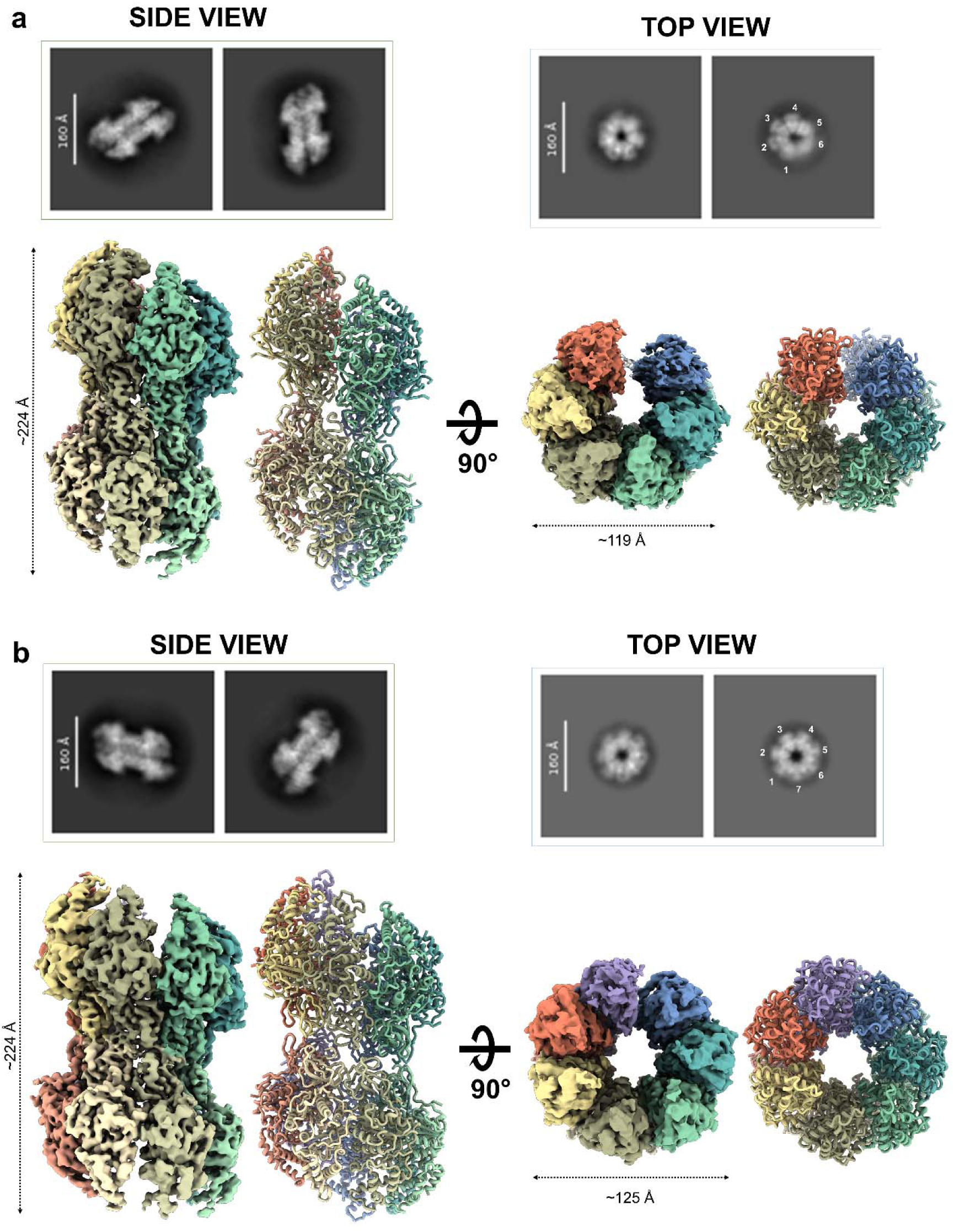
Structural overview of the dodecameric and tetradecameric YsaN ATPase assemblies. **a**, Top: Representative 2D class averages of the dodecameric (12-mer) YsaN assembly in orthogonal side and top views. Subunits in the top view are numbered 1–6 to indicate the hexameric tier. Bottom: The corresponding cryo-EM density map and atomic model, shown in side and top views related by a 90° rotation. **b**, Top: Representative 2D class averages of the tetradecameric (14-mer) YsaN assembly. Subunits in the top view are numbered 1–7 to indicate the heptameric tier. Bottom: The corresponding cryo-EM density map and atomic model, displayed as in **a**. In both panels, individual protomers are uniquely coloured to illustrate the distinct subunit packing and the continuous helical split-ring architecture.

**Table 1:** Cryo-EM data collection and refinement and validation statistics.

|  | 11 mer | 12 mer | 13 mer | 14 mer |
| --- | --- | --- | --- | --- |
| **EMDB ID** | EMD-65882 | EMD-66223 | EMD-65920 | EMD-66331 |
| **PDB ID** | 9WD4 | 9WTO | 9WF0 | 9WX6 |
| **Data collection** |  |  |  |  |
| Microscope | Talos Arctica |  |  |  |
| Camera | Falcon 4 |  |  |  |
| Camera mode | Counting |  |  |  |
| Magnification (nominal) | ×150,000 |  |  |  |
| Voltage (kV) | 200 |  |  |  |
| Data acquisition software | EPU |  |  |  |
| Exposure navigation | AFIS |  |  |  |
| Electron exposure (e<sup>-</sup> per Å<sup>2</sup>) | 40 |  |  |  |
| Defocus range (μm) | −0.5 to −2.0 |  |  |  |
| Pixel size (Å per pixel) | 0.95 |  |  |  |
| Micrographs collected (no.) | 14159 |  |  |  |
| Data analysis |  |  |  |  |
| Total extracted picks (no.) | 2760823 |  |  |  |
| Particles used for 3D (no.) | 709628 |  |  |  |
| Final particles (no.) | 174701 |  | 123350 |  |
| Symmetry imposed | C1 |  |  |  |
| Map resolution (global) (Å) | 3.51 |  | 3.80 |  |
| FSC□=□0.5 (unmasked/masked) | 7.24/3.97 |  | 7.56/4.19 |  |
| FSC□=□0.143 (unmasked/masked) | 4.11/3.51 |  | 4.27/3.80 |  |
| Local resolution range (Å) | 2.5–7.5 |  | 2.5–7.5 |  |
| Map sharpening B factor (Å²) | 54.0 |  | 50.3 |  |
| Model composition |  |  |  |  |
| Chains | 11 | 12 | 13 | 14 |
| Nonhydrogen atoms | 32702 | 37439 | 40780 | 44282 |
| Protein residues | 4332/4796 | 4993/5232 | 5255/5590 | 5698/6020 |
| Model refinement |  |  |  |  |
| Initial models used (AlphaFold Id) | AF-A1JQ97-F1 |  |  |  |
| Refinement package | UCSF ChimeraX, Coot, PHENIX |  |  |  |
| Bond lengths (Å) | 0.004 | 0.002 | 0.003 | 0.002 |
| Bond angles (°) | 0.516 | 0.576 | 0.640 | 0.514 |
| Validation |  |  |  |  |
| Ramachandran plot |  |  |  |  |
| Favored (%) | 95.84 | 96.84 | 97.87 | 98.53 |
| Allowed (%) | 4.06 | 3.12 | 2.09 | 1.47 |
| Outliers (%) | 0.09 | 0.04 | 0.04 | 0.00 |
| MolProbity score |  |  |  |  |
| Clashscore | 5.06 | 4.82 | 8.31 | 8.52 |
| Poor rotamers (%) | 0.69 | 1.20 | 0.97 | 1.19 |
| Q score | 0.3780 | 0.4050 | 0.3590 | 0.3160 |

A prominent feature of both oligomeric states is a longitudinal fissure on the cylindrical face of the assembly that breaks the radial symmetry. The local resolution estimate indicates that the protomers immediately adjacent to this fissure have lower density, reflecting intrinsic structural flexibility or dynamic motion in this region. During initial global refinement, the extreme flexibility of one protomer at the fissure resulted in maps where only 11 and 13 subunits were clearly resolved for the dodecameric and tetradecameric assemblies, respectively (Table 1). In contrast, protomers located on the opposite side of the ring display higher local resolution, suggesting a highly rigid and stable inter-subunit arrangement. To address the poorly resolved fissure protomers and generate complete density maps suitable for robust atomic modelling, we performed focused local refinement and merged these local maps with the global reconstructions (Fig. S2).

The full-length YsaN monomer is organized into three structurally distinct domains (Fig. 4a). The N-terminal oligomerization domain (residues 1–78) comprises a cluster of curved β-sheets. The central ATPase domain (residues 79–354) adopts a canonical Rossmann fold^43^ with nine parallel β-strands flanked by four α-helices on one side and three on the other. Finally, the C-terminal domain (residues 355–430) is composed of four α-helices that line the central pore of the oligomeric assembly. The central ATPase domain contains the highly conserved Walker A or P-loop^44^ (residues 160–167; AAAGSGKT) and Walker B^44^ (residues 246–251; LLFVDS) motifs, supporting its role in ATP binding and hydrolysis (Fig. 4a, Fig. S4). The overall folds of the ATPase and C-terminal domains are highly conserved among YsaN orthologues and related T3SS ATPases. In contrast, the N-terminal region, for which complete structural information is lacking, displays considerable sequence and structural variability (Fig. S4, 5). In this study, we present the full-length high-resolution cryo-EM structure of YsaN and reveal how its variable N-terminal domain shapes inter-subunit interactions and directly drives the assembly of higher-order cylindrical architectures.

### Helical geometry and longitudinal fissure dynamics

Both 12mer and 14mer have formed hollow cylindrical structures with a sharp fissure or opening longitudinally on the curved surface of the cylinder. In our calculation, we found that each protomer has an average rotation of ∼57–59° and a vertical translation of ∼6–7 Å along the central axis relative to its adjacent protomer, forming the lower part of the cylinder comprising six protomers. In the upper cylindrical part, the similar protomer arrangement continues, but the protomer vertically toggles so that the upper and lower cylinders are stabilized by the N-terminal regions of the protomers. As a result, the central part of the assembly is slightly tapered in appearance and constricts the inner diameter of the cylinder (Figure 2c). A little distortion appears in the vertical direction of the cylinder due to the ∼6–7 Å vertical translation of each protomer. The tetradecameric complex displays a similar structural architecture, with an average twist of ∼50–57° and a vertical translation of ∼5–7 Å per protomer (Figure 2c). This continuous helical progression is further stabilized by a conserved inter-subunit salt bridge. In our analysis of the 12-mer and 14-mer assemblies, we identified a conserved salt bridge between Arg192 and Asp324 of adjacent subunits. This interaction, located at the active-site interface, may also help stabilize the active-site architecture. Similarly, a conserved salt bridge between Arg209 and Asp340 is present in the EscN hexamer structure. Both the arginine and aspartate residues involved are highly conserved among T3SS ATPases. Notably, mutation of the analogous arginine residue in the Shigella T3SS ATPase Spa47 (Arg191) was previously shown to completely abolish oligomer formation^45^, underscoring the critical role of this conserved inter-subunit interaction in oligomer assembly. Collectively, these findings suggest that a network of conserved salt-bridge interactions contributes to T3SS ATPase oligomerization and may represent a universal target for disrupting oligomeric assembly in future structure-guided studies. This steady structural progression is sharply interrupted at the longitudinal fissure or opening, a pronounced structural discontinuity that seems to open the overall architecture. The dimensions of this fissure vary distinctly between the two oligomeric states. The vertical translation in the fissure region is almost similar, ∼35 Å in the 12mer and ∼38 Å in the 14mer (Fig. 2b, c, Movie S2), but the angular displacement in this region differs between the 12mer and 14mer. In the 12mer, the rotational twist across the fissure expands substantially to ∼71–74° (Fig. 2a). In contrast, the 14mer fissure is significantly compressed, exhibiting a reduced angular gap of ∼37–40° (Fig. 2a).

**Fig. 2.**
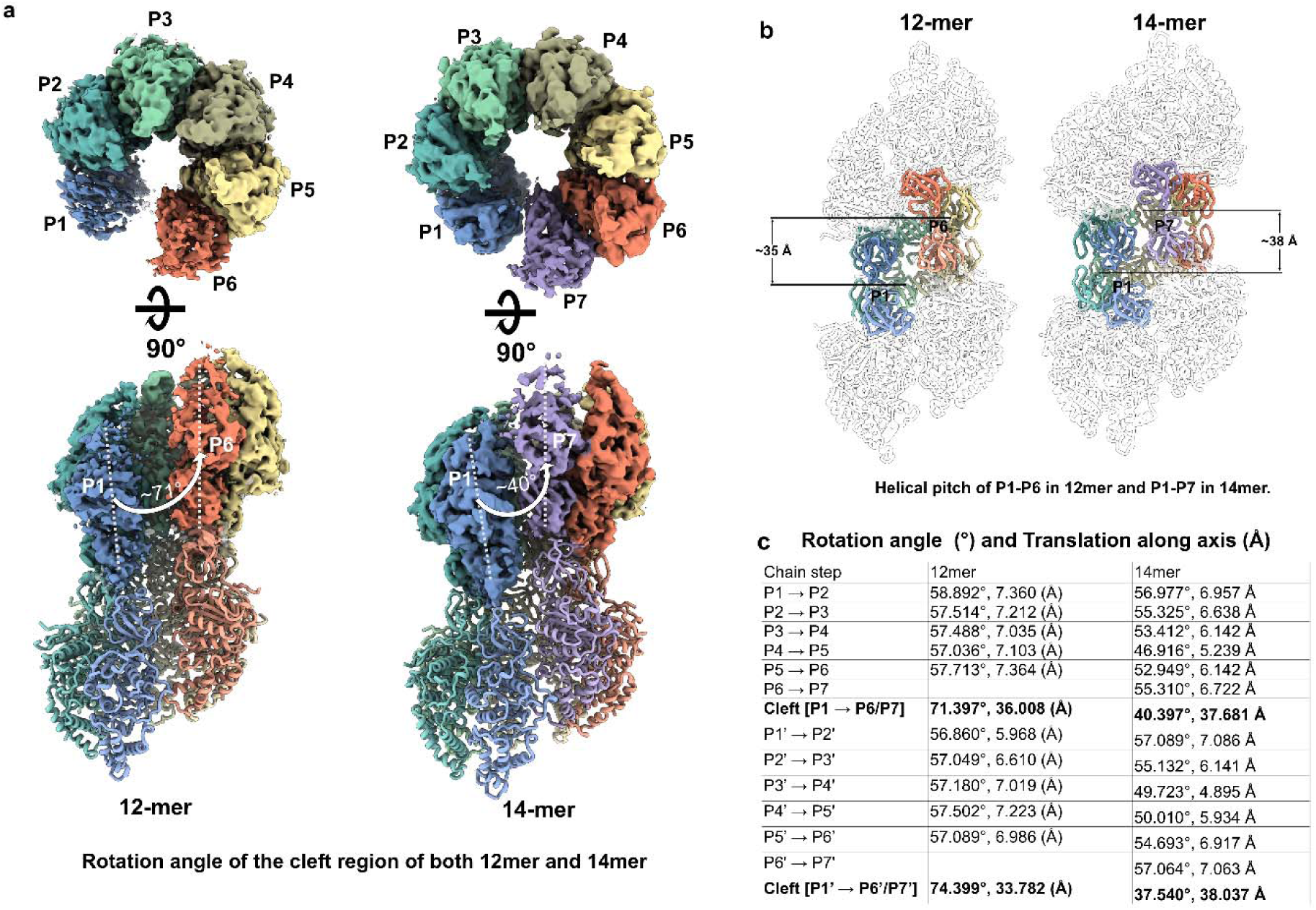
Helical geometry and asymmetric longitudinal fissure dynamics of the YsaN assemblies. **a,** Top and side views of the 12-mer (left) and 14-mer (right) fissure regions. In each panel, the top half displays the overall cryo-EM density map, while the bottom half displays the corresponding atomic model in the side view. Individual protomers (P1–P6 in the 12-mer; P1–P7 in the 14-mer) are distinctly coloured to trace the helical arrangement. The dashed white lines and arrows indicate the rotational twist across the longitudinal fissure, which expands to ∼71° in the dodecamer and is compressed to ∼40° in the tetradecamer. **b,** Structural outlines of the 12-mer and 14-mer architectures, where only the N-terminal domains are coloured to track the helical pitch. The solid black lines denote the massive vertical translational jump across the fissure (∼35 Å and ∼38 Å, respectively), highlighting the pronounced physical opening of the split-ring geometry. **c,** Tabulated rotation angles and vertical translations between adjacent protomers along the central helical axis. The values demonstrate a steady helical progression that is sharply interrupted by massive structural discontinuities at the dynamic fissure interface (bolded), underscoring the stoichiometric plasticity between the two states.

The massive angular gap observed in the dodecameric fissure provides sufficient physical space to accommodate a pair of additional subunits. This strongly suggests that the longitudinal fissure serves as a dynamic docking site, enabling the complex to structurally interconvert between the 12-mer and 14-mer states, expanding or contracting the central tunnel based on specific physiological or substrate-handling requirements.

### N-terminal domains mediate oligomeric assembly

The integrity of this helical architecture is fundamentally driven by an extensive non-covalent and covalent interaction network between adjacent and opposing subunits. Interface analysis revealed that adjacent protomers bury an average surface area of ∼1,230 Å² and are stabilized by hydrogen bonds and inter-subunit salt bridges. Crucially, the dense helical packing firmly locks the upper and lower tiers of the complex together, burying a substantial inter-tier surface area of 1,229.4 Å² in the 12-mer and 1,258.3 Å² in the 14-mer. These vital inter-tier contacts are primarily mediated by the extreme N-terminal residues, a region that has been truncated in all previously reported T3SS ATPase structures. Within these tightly packed inter-tier interfaces, we observed continuous cryo-EM density corresponding to six and seven disulfide bonds in the 12-mer and 14-mer, respectively, forming between closely apposed Cys38 residues on opposite protomers (Fig. 3a, b). Notably, sequence alignment reveals that Cys38 is a non-conserved residue among homologous T3SS ATPases, suggesting a unique evolutionary adaptation in YsaN (Fig. 3c).

**Fig. 3.**
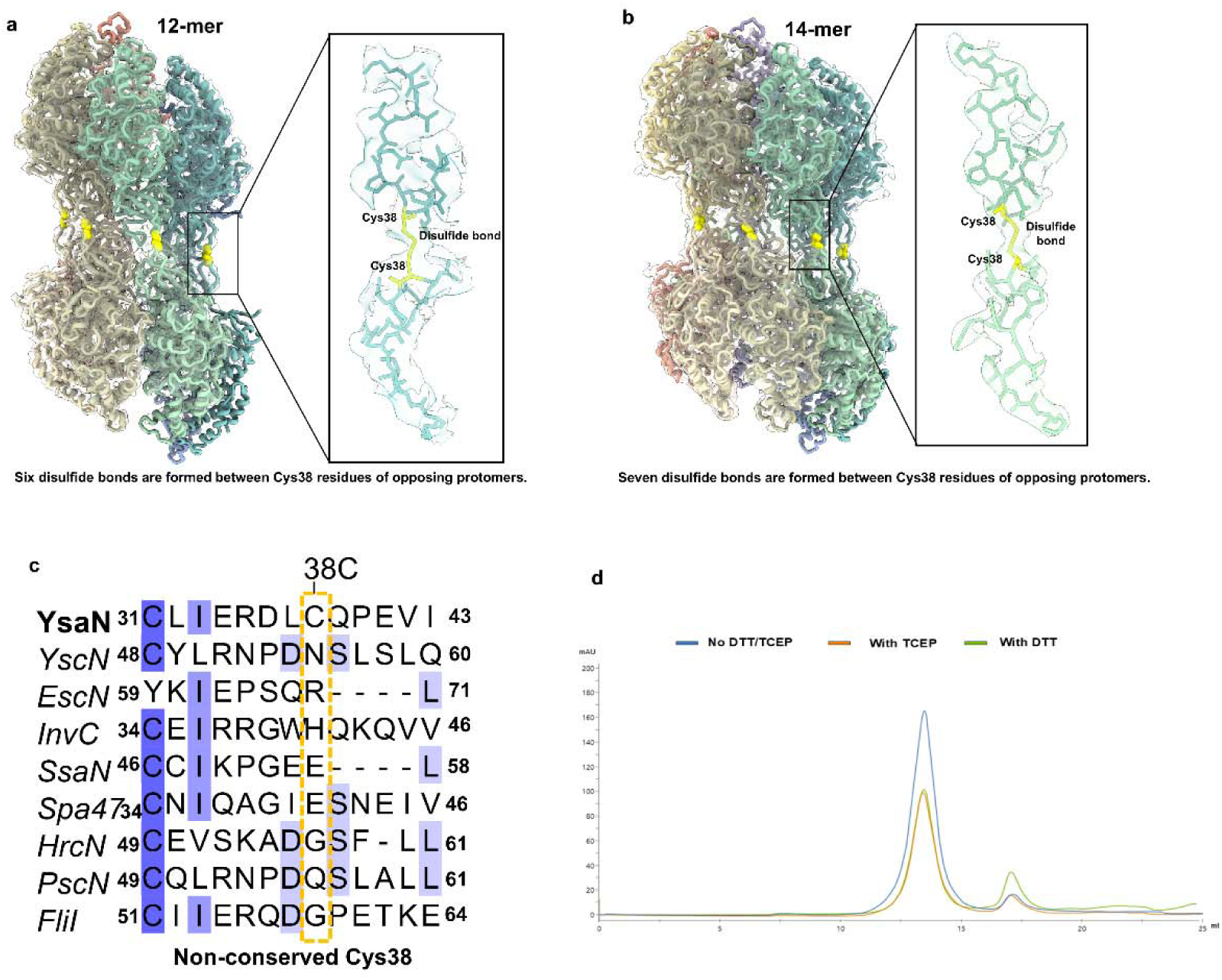
The N-terminal domain mediates the double-tiered helical assembly of YsaN. **a,** Side view of the dodecameric (12-mer) YsaN assembly, with the atomic model fitted inside the transparent cryo-EM density. Yellow spheres highlight the locations of the six disulfide bonds formed between the upper and lower tiers. The magnified inset details one representative disulfide bond between two opposing protomers, displayed as a stick model with the Cys38 linkage (yellow) clearly resolved within the transparent cryo-EM density. **b,** Corresponding side view of the tetradecameric (14-mer) assembly. **c,** Multiple sequence alignment of the extreme N-terminal region of YsaN alongside homologous T3SS and flagellar ATPases. The alignment highlights that Cys38 (dashed yellow box) is a uniquely non-conserved residue, representing a specific evolutionary adaptation in YsaN for additional covalent structural reinforcement. **d,** Analytical size-exclusion chromatography (SEC) elution profiles of YsaN in the absence of reducing agents (blue line), or in the presence of 1 mM TCEP (orange line) or 5 mM DTT (green line). The primary peak corresponding to the intact higher-order oligomer remains structurally undisturbed upon reduction, demonstrating that the robust non-covalent interaction network of the N-terminus independently maintains the double-tiered assembly.

**Fig. 4.**
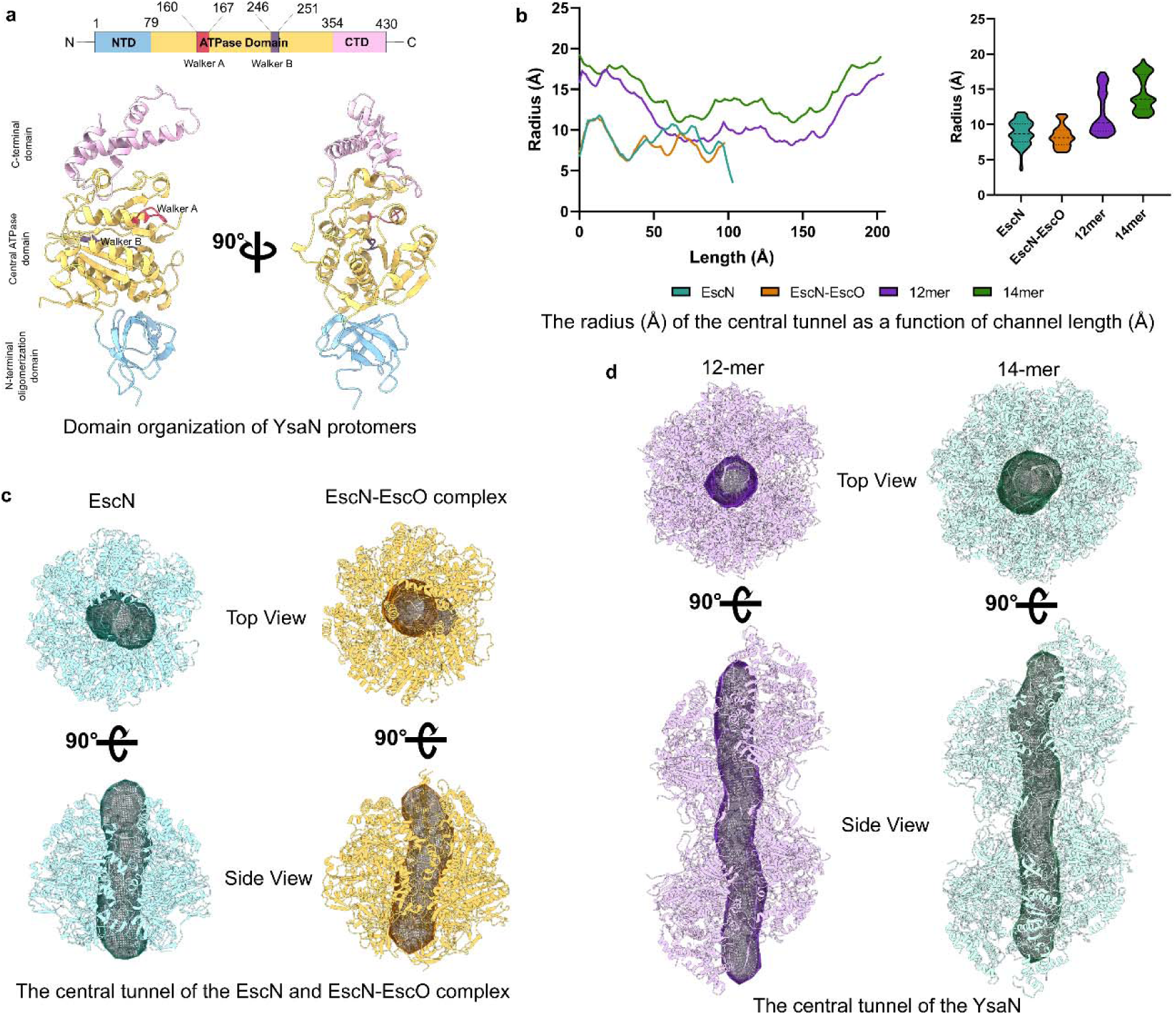
Protomer architecture and quantitative analysis of the central translocation cavity. **a,** Top: 1D domain organization of full-length YsaN highlighting the N-terminal domain (NTD, blue), central ATPase domain (yellow) containing the Walker A (red) and Walker B (purple) motifs, and the C-terminal domain (CTD, pink). Bottom: Ribbon representation of a single YsaN protomer shown in two orthogonal views. **b,** Left: CAVER analysis measuring the tunnel radius (Å) as a function of channel length (Å) for EscN (PDB id: 6NJO) (teal), EscN-EscO (PDB id: 6NJP) (orange), the YsaN 12-mer (purple), and the YsaN 14-mer (green). Right: Violin plots illustrating the overall distribution of pore radii across the length of each respective complex. **c,** Top and side views of the hexameric EscN and EscN-EscO structures, with the internal cavities calculated by CAVER represented as solid mesh surfaces. **d,** Top and side views of the expanded YsaN 12-mer and 14-mer helical assemblies, demonstrating the significant global radial expansion of the central translocation tunnels.

While these covalent linkages provide an additional layer of structural stabilization to the mature complex, we sought to determine if they were the sole obligate requirement for holding the two tiers together. We performed size-exclusion chromatography (SEC) in the presence of reducing agents, 5 mM DTT and also with 1 mM TCEP. Strikingly, the SEC elution profiles of reduced YsaN remained essentially identical to those under non-reducing conditions, eluting predominantly as intact higher-order oligomers and monomers with no dissociation into hexameric or heptameric halves (Fig. 3d). This demonstrates that buried surface area is driven uniquely by the intact N-terminus, which is sufficiently robust to independently maintain the double-tiered helical assembly, with the Cys38 disulfide bonds serving to further reinforce the structural integrity of the complex.

### Oligomeric state dictates the geometry of the central translocation tunnel

To quantitatively assess how stoichiometric plasticity influences the geometry of the internal secretion conduit, we analysed the central translocation tunnels of the dodecameric and tetradecameric assemblies using CAVER^46^ (Fig. 4b–d). Both oligomeric states display a continuous central passage; however, the tetradecamer exhibits a markedly wider pore across its entire longitudinal axis. The minimum radius (the bottleneck region) in the dodecamer is approximately 8–9 Å (corresponding to a diameter of 16–18 Å). In contrast, the tetradecamer displays a substantially expanded bottleneck radius of 11–12 Å (22–24 Å diameter). Beyond this constriction point, the pores progressively widen toward both exits, reaching maximal diameters of 30–35 Å in the dodecamer and 36–38 Å in the tetradecamer (Fig. 4b). The hexameric T3SS ATPase EscN also forms a central tunnel featuring a diameter of ∼15-25 Å (Fig. 4c).

These observations indicate that these assemblies feature central tunnels large enough to accommodate protein substrates in a semi-unfolded state and sufficiently wide to allow the passage of bulky, semi-folded structural elements.^47,48^

### ATPase activity is regulated by oligomerization

Consistent with our previous study, in this study we observed that the higher-order oligomeric fraction (comprising the 12-mer and 14-mer assemblies) exhibited higher ATPase activity that quickly reached maximum substrate turnover. In contrast, the monomeric YsaN fraction displayed negligible catalytic activity (Fig. 5a). This confirms that, similar to previous reports for homologous systems, the assembly of YsaN into higher-order ring architectures is needed for its enzymatic function^29,31,38,41,49^.

**Fig. 5.**
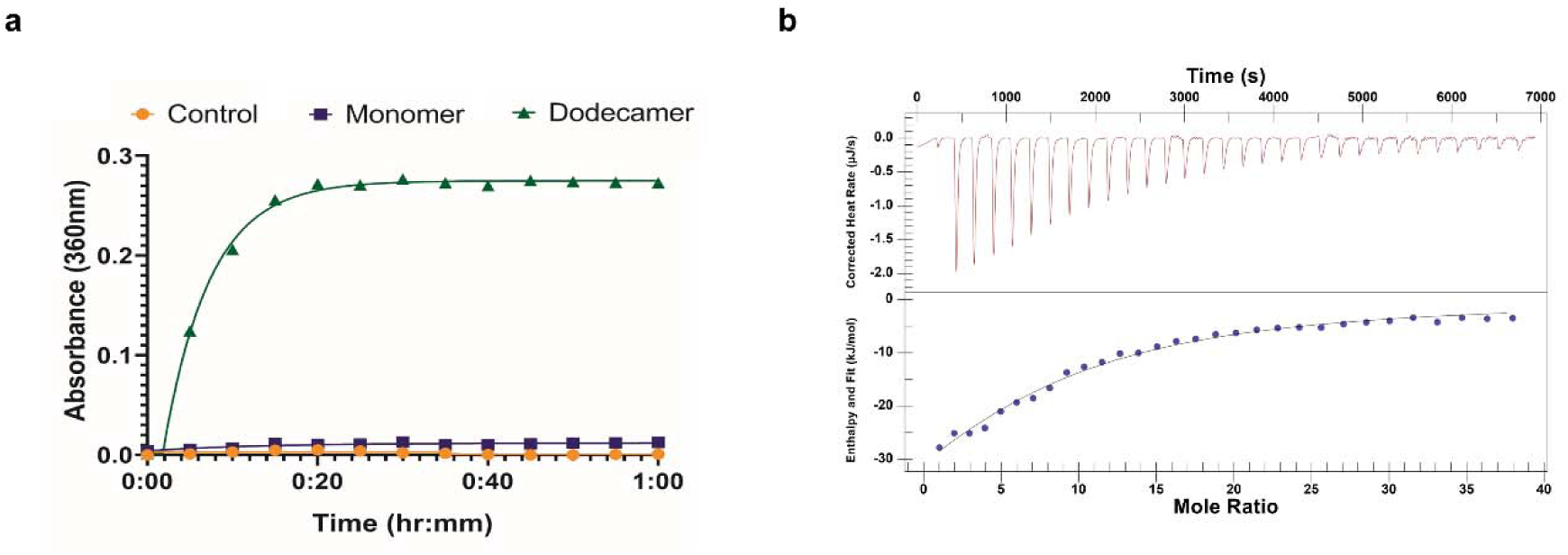
Oligomerization is required for YsaN ATPase activity and nucleotide engagement. **a,** ATP hydrolysis kinetics measured via a phosphate release assay. The higher-order oligomeric YsaN fraction (green triangles) exhibits rapid catalytic activity that quickly reaches maximum substrate turnover. In contrast, the monomeric fraction (purple squares) displays very little ATPase activity, the baseline buffer control shown as orange circles. This confirms that higher-order assembly is an absolute prerequisite for YsaN enzymatic function. **b,** Representative isothermal titration calorimetry (ITC) thermogram (top) and corresponding integrated binding isotherm (bottom) for the higher-order YsaN complex titrated with ATPγs. The integrated heat data fit a single-site binding model.

To further study binding of nucleotides to the active assemblies of the protein, we performed isothermal titration calorimetry (ITC) using ADP and the non-hydrolysable ATP analogue, ATPγS. The nucleotide-binding events were exothermic, and the integrated ITC data were best fit by a single-site binding model (Fig. 5b). The thermodynamic parameters obtained from these calculations are listed in Table 2. ITC experiments showed that the dissociation constant (K_d_) of YsaN for ATPγS and ADP are 461.9 ± 79.2 µM and 383.3 ± 57.2 µM, respectively. Crucially, the measured binding stoichiometries (n) ≈ 9.3 for ATPγS and 10 for ADP, align closely with our structural characterization of YsaN as a 12-mer/14-mer complex. This confirms that nearly every protomer within the higher-order ring is capable of actively engaging a nucleotide in a concerted manner. The entropic parameters indicate that the ADP-bound state possesses lower structural entropy compared to the ATPγS-bound state (ADP: ΔS = -287.8 J mol⁻¹ K⁻¹; ATPγS: ΔS = -143.5 J mol⁻¹ K⁻¹).

**Table 2:** Thermodynamic Parameters for The Interaction of ATPγS and ADP With YsaN. Thermodynamic parameters were obtained from an average of three independent ITC experiments.

| Protein-Ligand | T(K) | n | $K_d \times 10^{-4} \text{ M}^{-1}$ | $\Delta G \text{ kJ mol}^{-1}$ | $\Delta H \text{ kJ mol}^{-1}$ | $\Delta S \text{ J mol}^{-1} \text{ K}^{-1}$ |
| --- | --- | --- | --- | --- | --- | --- |
| YsaN-ATPyS | 283 | $9.3 \pm 1.6$ | $4.6 \pm 0.8$ | -20.9 | $-61.5 \pm 17.9$ | -143.5 |
| YsaN-ADP | 283 | $10.0 \pm 0.5$ | $3.8 \pm 0.6$ | -18.5 | $-100.0 \pm 7.3$ | -287.8 |

### Molecular dynamics analysis reveals nucleotide-dependent conformational changes of YsaN oligomeric assemblies

Given that YsaN is the central energetic driver of the *Yersinia* T3SS, ATP binding and hydrolysis are expected to play a critical role in modulating its conformational stability, oligomeric organization, and catalytic architecture. To investigate the structural consequences of nucleotide engagement, we performed 200 ns all-atom molecular dynamics (MD) simulations of the dodecameric (12-mer) and tetradecameric (14-mer) YsaN assemblies in their apo, ADP-bound, and ATP-bound states. Interaction mapping of the nucleotide-bound models revealed that the nucleotides are predominantly coordinated by residues within the highly conserved Walker A motif (residues 160–167; AAAGSGKT) (Fig. S6).

Specifically, Ser164, Gly165, Lys166, and Thr167 form extensive hydrogen-bonding, electrostatic, and water-mediated interactions with the phosphate groups. Notably, Lys166 establishes stabilizing contacts with the β/γ-phosphate region, consistent with its conserved role as the primary phosphate-binding lysine in P-loop ATPases, while Tyr339 forms a stacking interaction with the adenosine base.

To evaluate the global effect of nucleotide binding on oligomeric stability and conformational dynamics, we analysed the trajectories of the 200 ns MD simulations (Movies S3, S4). Root-mean-square deviation (RMSD) analysis indicated that all systems reached equilibrium after approximately 50 ns and remained stable for the remainder of the simulation (Fig. 6a); accordingly, the final 150 ns of the trajectories were utilized for detailed structural analysis. While the mean RMSD values are broadly comparable between the two oligomeric states, the density distributions reveal that the 14-mer assemblies exhibit broader structural deviations, indicative of greater intrinsic flexibility compared to the 12-mer (Fig. 6a). Root-mean-square fluctuation (RMSF) profiles are highly similar between the 12-mer and 14-mer, demonstrating that the C-terminal domain is substantially more flexible than the N-terminal oligomerization and central ATPase domains, which perfectly mirrors the local resolution distribution observed in our cryo-EM maps (Fig. 6c). Furthermore, the RMSF profiles confirm that the ADP-bound state represents the most dynamically stable and rigid conformation for both oligomeric assemblies. This observation is consistent with our ITC measurements, which showed a lower entropy for ADP-bound YsaN than for the ATPγS-bound state, suggesting reduced conformational flexibility upon ADP binding.

**Fig. 6.**
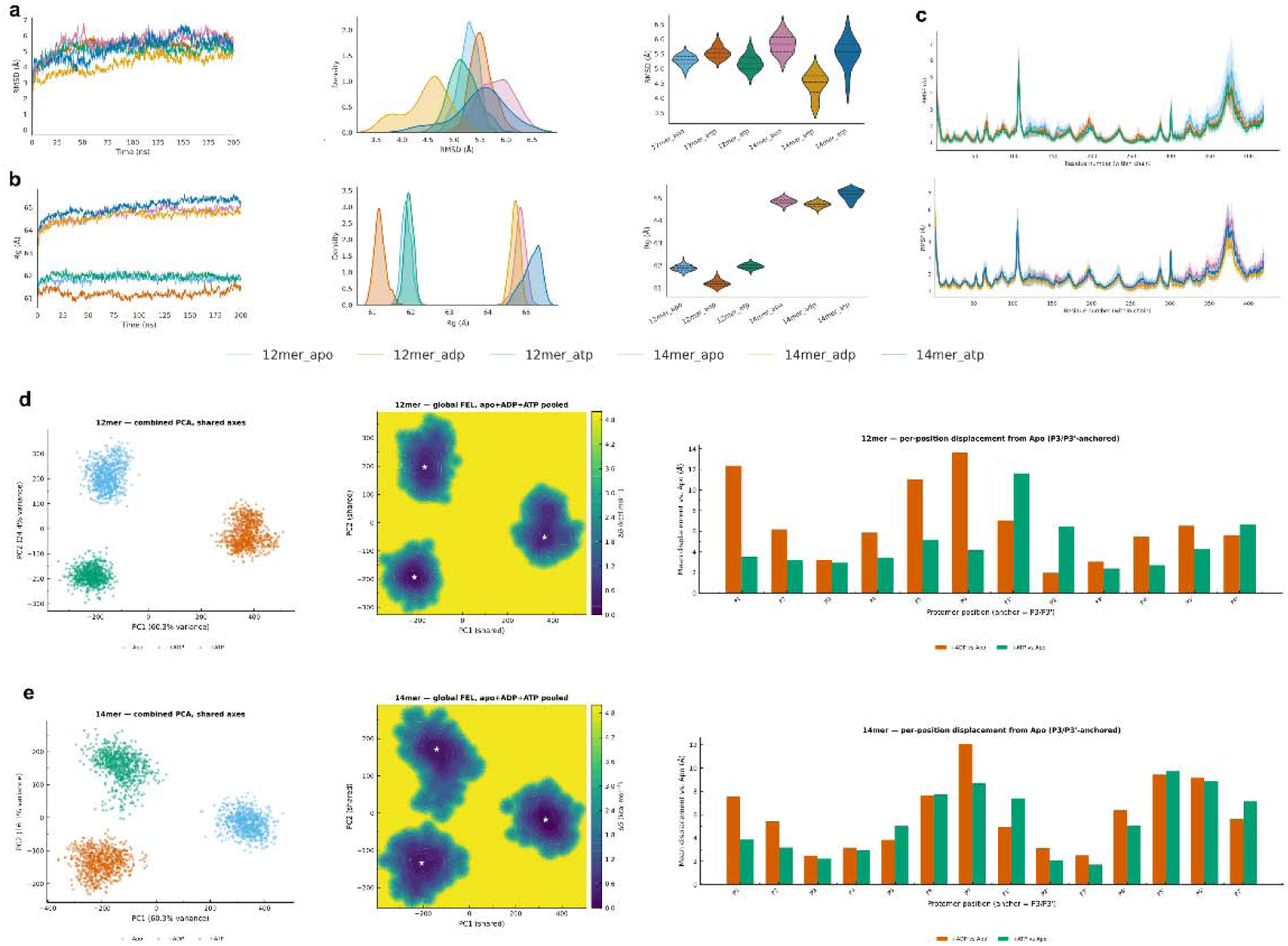
Molecular dynamics simulations reveal a nucleotide-driven global compaction and longitudinal fissure-closing power stroke. **a,** Root-mean-square deviation (RMSD) analysis of the 12-mer and 14-mer YsaN assemblies over 200 ns simulations, displaying the time-dependent trajectories, density distributions, and violin plots. The data indicate that all systems reach a stable equilibrium, with the 14-mer exhibiting slightly broader structural deviations. **b,** Radius of gyration (Rg) analysis illustrating global structural compaction upon nucleotide binding. The trajectory tracking, density plots, and violin plots demonstrate that the ADP-bound state adopts a significantly tighter, more compact architectural footprint compared to the apo and ATP-bound states. **c,** Root-mean-square fluctuation (RMSF) profiles per residue for the 12-mer (top) and 14-mer (bottom) assemblies, confirming that the C-terminal domains exhibit the highest intrinsic flexibility. **d, e,** Conformational and mechanical landscape of the 12-mer (**d**) and 14-mer (**e**) assemblies. Left: Principal component analysis (PCA) projections (PC1 vs. PC2) show that the apo, ADP-bound, and ATP-bound states occupy entirely distinct conformational spaces. Middle: Free energy landscapes (FEL) confirm that each nucleotide state settles into a well-isolated thermodynamic basin (global minima indicated by white stars). Right: Per-position structural displacement analysis anchored at the rigid opposing protomers (P3/P3’). The bar graphs reveal extreme functional asymmetry, demonstrating that the most massive nucleotide-driven structural rearrangements occur almost exclusively at the protomers directly flanking the longitudinal fissure (P1 and P6 in the 12-mer; P1 and P7 in the 14-mer).

Tracking the radius of gyration (Rg) over the simulation trajectories highlighted a nucleotide-driven structural compaction. In both the 12-mer and 14-mer assemblies, the ADP-bound state exhibits a significantly smaller Rg compared to the apo and ATP-bound structures (Fig. 6b). This demonstrates that ATP hydrolysis and the subsequent transition to the ADP-bound state induce a global compaction.

To map the thermodynamic transitions governing this mechanochemical cycle, we performed principal component analysis (PCA) and generated free energy landscapes (FEL) for the simulated states (Fig. 6d, e). Projection of the trajectories onto the first two principal components (PC1 and PC2) revealed that the apo, ATP-bound, and ADP-bound states occupy entirely distinct, non-overlapping conformational spaces. The FEL confirms that each nucleotide state settles into a well-defined, isolated thermodynamic basin.

To quantify the physical mechanics of these state transitions, we measured the per-protomer structural displacement relative to the apo state, anchoring our analysis at the rigid protomers directly opposite the fissure (P3/P3’). The displacement analysis shows functional asymmetry: the most structural rearrangements occur almost exclusively at the protomers directly flanking the longitudinal fissure (P1/P1’ and P6/P6’ in the 12-mer; P1/P1’ and P7/P7’ in the 14-mer) (Fig. 6d, e). As visualized by the dominant principal motions extracted from the trajectories (Movies S5-7), nucleotide turnover drives a synchronized, global twisting motion that culminates in the physical closing of the longitudinal fissure. Together, these data demonstrate that YsaN operates as a highly dynamic machine, where the chemical energy of ATP hydrolysis is translated into a synchronized global compaction and fissure-closing “power stroke” to drive substrate threading.

### The conserved aromatic stacking residue Tyr339 occludes the adenine binding hydrophobic pocket in YsaN

In the ligand-bound structures of EscN (PDB ID: 6NJO) and InvC (PDB ID: 6SDX), the adenine moiety is accommodated within a hydrophobic pocket and further stabilized by π–π stacking with a conserved aromatic residue. A comparable hydrophobic pocket is present in YsaN, lined by Leu329, Leu169, Pro340, Ala341, Met172, and Ile342. With the exception of Met172, these residues are fully conserved across T3SS ATPases from diverse species (Fig. S7c). In YsaN, however, the pocket is occluded by the side chain of the conserved adenine-stacking residue Tyr339 (Fig. S7a). Superposition of all six EscN subunits with YsaN chains revealed that the Tyr339 side chain adopts a ∼90° rotated conformation relative to Phe355 in EscN, consistently across all YsaN subunits (Fig. S7d). This observation suggests that ligand binding may induce a local rearrangement of Tyr339, exposing the hydrophobic pocket and accommodating the adenine moiety. The assumption is further validated by examining the ATP-docked structure of YsaN 12-mer, which shows opening of the pocket and movement of Tyr339 side-chain to accommodate the adenine in the hydrophobic pocket (Fig. S7b).

A similar mechanism of pocket opening is evident from comparison of the apo and ligand-bound structures of the Salmonella ATPase InvC (PDB IDs: 6RAE and 6SDX, respectively). Although the adenine-stacking aromatic residue adopts essentially the same conformation in both states of InvC, a methionine (Met167) side chain occludes the pocket in the InvC apo structure and is moved upon ligand binding (Bernal et al. 2019). Together, these structural comparisons and in silico evidence suggest that local side-chain rearrangements may regulate access to the conserved adenine-binding pocket in T3SS ATPases, thereby enabling accommodation of the adenine moiety upon nucleotide binding.

## Discussion

T3SS ATPases are central to substrate recognition, unfolding, and translocation^5^, yet the higher-order assembly and mechanistic organisation of these enzymes remain incompletely understood. Structural studies to date have focused largely on the hexameric assembly^31,26,28^. Our cryo-EM analysis of full-length YsaN instead reveals two distinct higher-order oligomeric states, a dodecamer and a tetradecamer.

Dodecameric assembly has been proposed previously^38,41^, including in our own earlier work, but has not until now been visualised directly. Here we report both the dodecamer and the tetradecamer at 3.5 Å and 3.8 Å resolution, respectively, sufficient to resolve the molecular determinants of their assembly and to propose a mechanistic model for substrate translocation.

Assembly of both oligomers is maintained principally by the N-terminal oligomerisation domain, a region that is partially or completely absent from all previously reported homologue structures^26,29,31,33^. This domain is helically arranged and couples the upper and lower tiers of the complex, with the resulting inter-tier interface burying 1,229.4 Å² of surface area in the dodecamer and 1,258.3 Å² in the tetradecamer. At the core of each interface, we identified continuous density consistent with disulfide bridges, six in the dodecamer and seven in the tetradecamer, linking Cys38 residues from opposite protomers. Because

Cys38 is not conserved across other T3SS ATPases, we suggest that this disulfide bonds represents a distinct, potentially YsaN-specific mode of stabilising the oligomeric interface. This indicates that the N-terminal domain is a key determinant of higher-order oligomer formation in YsaN. Adjacent protomers are further stabilised by a network of hydrogen bonds and salt bridges, an interaction mode consistent with the dimeric and trimeric species reported in several earlier biochemical studies^26,28,30^.

Consistent with these structural findings, the higher-order oligomers displayed substantially greater ATPase activity than the monomer, which showed minimal activity, an observation in agreement with our previous detailed kinetic characterisation and with reports for other T3SS ATPases^28,38,41^. Neither oligomer possesses overall symmetry; instead, each displays a fissure, which is also seen in EscN^31^. EscN itself likewise lacks strict symmetry, but its protomers are nonetheless arranged in a broadly cyclic manner, whereas the YsaN oligomers instead adopt a distinctly helical arrangement. Building on the established model in which substrate–chaperone complexes dock at the C-terminal domains near the fissure, with ATP hydrolysis driving dissociation of the substrate from its chaperone and its consequent unfolding to a semi-unfolded state^21,28,47,50^, we propose an additional step specific to this helical architecture: the semi-unfolded substrate then enters through the fissure into the central tunnel of the YsaN oligomer, from where it is transported to the export apparatus via a threading-like mechanism (Fig. 7). Measurement of the central pore in both assemblies, relative to the EscN hexamer, revealed a marked expansion of the constriction point: the dodecamer presents a minimum bottleneck diameter of 16–18 Å, whereas the tetradecamer is substantially wider at this position, at 22–24 Å. Beyond this constriction, the pore widens progressively toward both ends of the channel, reaching maximal diameters of 30–35 Å in the dodecamer and 36–38 Å in the tetradecamer, dimensions sufficient to accommodate the substrate in its unfolded state^51,52^.

**Fig. 7.**
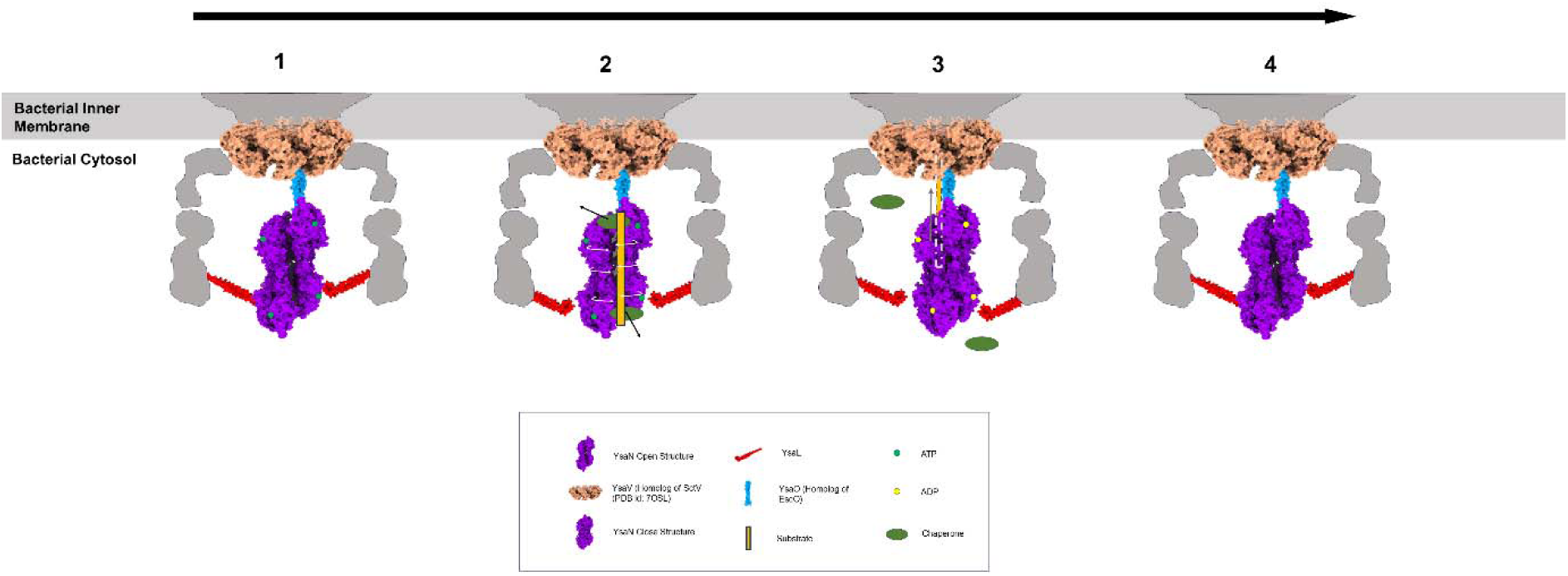
Proposed mechanochemical threading model of the YsaN ATPase. **1,** In the apo or ATP-bound state (green spheres), the YsaN assembly (magenta) adopts an open, highly flexible conformation within the bacterial cytosol, positioned beneath the central stalk (YsaO, blue) and the inner membrane export apparatus (YsaV, peach). **2,** The intrinsic structural flexibility and widened central pore of the open state create a receptive environment for the docking of the bulky chaperone-substrate complex (green oval and yellow bar, respectively). **3,** ATP hydrolysis to ADP (yellow spheres) triggers a synchronized global structural compaction of the YsaN complex into a rigid, closed conformation (dark purple). This nucleotide-driven mechanical power stroke facilitates chaperone stripping and forcibly threads the unfolded substrate upward through the pore toward the export gate. **4,** Following successful substrate translocation, the complex resets to the open, substrate-receptive conformation to initiate the next cycle

We also examined the geometry of the inter-protomer fissure in each state. The angular discontinuity across this fissure differs markedly between the two oligomers: in the dodecamer, the rotational twist widens to approximately 71–74°, whereas in the tetradecamer it is compressed to approximately 37–38°. The pronounced angular gap at the dodecameric fissure provides sufficient space to accommodate an additional pair of subunits, raising the possibility that this fissure acts as a dynamic docking site through which the complex interconverts between the 12-mer and 14-mer states, thereby expanding or contracting the central translocation chamber according to physiological or substrate-handling demands.

This interpretation is supported by molecular dynamics simulations, in which the ADP-bound structure was consistently more compact and more stable than either the ATP-bound or apo forms, with a correspondingly narrower fissure. We propose that this closed, ADP-bound conformation limits substrate egress through the fissure and instead favours substrate movement toward the export-gate apparatus. Isothermal titration calorimetry (ITC) measurements showed that both the ADP and ATP binds very tightly with YsaN, consistent with previously reported data^28^. The ITC data also shows that upon ADP binding the protein gets structurally compact and stable as evident from decrease in entropy, in agreement with the MD results. Local resolution in both cryo-EM maps was reduced specifically at the protomers flanking the fissure, a feature also reported for EscN, indicating conformational mobility at this interface consistent with lid-like opening and closing. The C-terminal domain likewise showed lower local resolution than either the ATPase domain or the N-terminal domain, suggesting inherent flexibility that may facilitate its proposed role in detaching substrate from the chaperone.

A principal limitation of this study is that the involvement of YsaO in the assembly and translocation mechanism remains undefined. Future work will address this by determining cryo-EM structures of YsaN in complex with YsaO in the nucleotide bound forms.

Taken together, these findings position YsaN as a structurally dynamic ATPase whose oligomeric state is directly coupled to its transport function. Rather than existing as a single fixed assembly, YsaN interconverts between a dodecameric and a tetradecameric form through changes in fissure geometry, with nucleotide state acting as a switch that stabilises a compact, ADP-bound conformation favouring directional substrate movement. The N-terminal oligomerisation domain, largely unresolved in prior homologue structures, emerges as the structural determinant that permits this higher-order assembly in the first place. Together with the biochemical and computational data presented here, these structures extend the current hexamer-centric view of T3SS ATPases and support a model in which oligomeric plasticity, not a fixed ring architecture, underlies substrate unfolding and translocation. Resolving how YsaO integrates into this cycle will be an important next step toward a complete mechanistic picture of the export process.

## Methods

### Protein purification

Recombinant YsaN was expressed in Escherichia coli BL21 DE3 cells following a previously established protocol^38,53^. Briefly, a secondary culture was grown at 37 °C to an OD₆₀₀ of ∼0.6, followed by induction with 0.75 mM IPTG and incubation overnight at 25 °C to promote protein overexpression. Cells were harvested by centrifugation at 6000× g for 10 min, and pellets were lysed or stored at −80°C for future use. The pellet was resuspended in buffer A containing 50 mM Tris-HCl (pH 8.0), 150 mM NaCl, and 5 mM MgCl₂. The cells were lysed by sonication, and the clarified lysate was applied to a Ni²⁺-NTA affinity column for purification. The column was washed with buffer A supplemented with 25 mM imidazole to remove non-specifically bound proteins, and YsaN was eluted using buffer A containing 300 mM imidazole. The eluted protein was further purified by size-exclusion chromatography using a Superose 6 Increase column equilibrated in buffer A.

### ATPase assay

The ATPase assay was performed using the Invitrogen EnzChek^TM^ Phosphate Assay Kit (Invitrogen™) following its protocol. A protein concentration of 500 nM of monomer and 12mer/14mer was mixed in a reaction mixture containing 10 mM Tris, pH 7.5, 5 mM MgCl_2_, 0.5 mM ATP in the total volume of 200 μL in 96-well plates at 25 °C. After adding ATP to the reaction mixture, the absorbance of the secondary product was recorded at an interval of 5 mins for 60 mins using a microplate reader at 360 nm. The reaction sample without protein was taken as a blank for all experiments. All assays were performed in triplicate.

### Negative stain TEM

The sample was diluted to 25 µg/ml with buffer A. The 3.5 µL sample was applied for 10 s to carbon-coated 400 mesh Cu grids, which had been glow-discharged at 15 mA for 15 s (PELCO easiGlow™ Glow Discharge Cleaning System). The complex was negatively stained with 2% (w/v) uranyl acetate for 30 s. Data were collected on a Talos L120C Transmission Electron Microscope operating at 120 keV. Nominal magnification was 57,000 X with a pixel size of 2.536 Å at the specimen plane. The nominal defocus range was set around −1.5 µm. A fast-readout Ceta camera was used to record micrographs manually. The data were processed using cryoSPARC. Particles were selected from micrographs using blob picker. A total of ∼31,000 particles were extracted from 35 micrographs. Approximately ∼18,000 particles were selected using 2D classification.

### Sample preparation & data acquisition for Cryo-EM

The sample (3 µL) was applied at a concentration of ∼4.5 mg/mL and ∼3 mg/mL to Quantifoil R 1.2/1.3 300 Mesh, Copper Grid & Ultra Foil R 1.2/1.3 300 Mesh, Gold grid(Electron Microscopy Sciences), respectively, after being glow-discharged at 15 mA for 30 s & 60 s respectively (PELCO easiGlow™ Glow Discharge Cleaning System) and plunge-frozen into nitrogen-cooled liquid ethane using the Vitrobot mark IV (FEI). The settings for the Vitrobot were as follows: Temperature: 16 °C, humidity: 95%, blotting time: 2.5 s with standard Vitrobot filter paper, Ø55/20mm, Grade 595, blotting force: 2, and wait time: 5 s.

Cryo-EM data were collected over two sessions using a 200-kV Talos Arctica transmission electron microscope equipped with a Falcon 4 direct electron detector (Thermo Fisher Scientific). Automated acquisition was driven by EPU software (Thermo Fisher Scientific) utilizing aberration-free image shift (AFIS) mode to maximize acquisition speed and throughput. Micrographs were recorded as dose-fractionated movies in EER format at a nominal magnification of 150,000×, yielding a physical pixel size of 0.95 Å. The total dataset comprises 14,245 micrographs, each acquired with a total electron dose of 40 e−/Å² and a nominal defocus range of −0.5 to −2.5 µm.

### Cryo-EM data processing

Data processing was performed in cryoSPARC^54^ (version 4.6.2). To compensate for beam-induced motion and radiation damage, dose-fractionated movies (40 fractions, 40 e⁻/Å² total dose) were processed using the patch motion correction algorithm. Patch contrast transfer function (CTF) estimation was then applied to the aligned and summed micrographs. A curated subset of 10,633 good micrographs was retained for downstream analysis, selected based on minimal astigmatism and high-confidence CTF fits extending to at least 4 Å resolution.

To improve particle contrast, the micrographs were initially denoised. Blob picking followed by preliminary 2D classification was used to generate references for automated template picking. A total of 2,760,823 particles were extracted from the original, non-denoised micrographs using an 800-pixel box size, which was subsequently binned to 600 pixels (yielding a pixel size of 1.267 Å).

Iterative rounds of 2D classification, *ab initio* 3D reconstruction, and multiple iterations of heterogeneous refinement were employed to filter out poorly aligned particles and isolate the highest-quality structural subsets, and we identified two dominant structural states: a dodecameric complex consisting of two stacked hexamers, and a tetradecameric complex consisting of two stacked heptamers. Final 3D reconstructions were performed using Non-Uniform Refinement with 174,701 particles for the dodecamer and 123,350 particles for the tetradecamer. Based on the gold-standard Fourier shell correlation (FSC)^55^ threshold of 0.143, these maps achieved overall resolutions of 3.51 Å and 3.80 Å, respectively.

Because the initial consensus maps exhibited weak density for a single subunit in both assemblies, focused refinements were performed on these specific regions to recover the missing signal. The resulting focused maps were aligned and resampled to the grid space of the initial non-uniform refinement maps. Finally, comprehensive composite maps for both assemblies were generated in UCSF ChimeraX^56^ using the ‘volume maximum’ function, which retains the highest voxel values across the aligned volumes to complete the structures.

### Atomic model building into EM maps

To build the model of YsaN into the reconstruction, an initial model based on the Alphafold^57^ predicted model was used. The model was first rigid fitted into the map using UCSF ChimeraX, followed by iterative manual adjustment in COOT^58^, and then refined with phenix.real_space_refine^59^. Because certain protomers exhibited lower local resolution, non-crystallographic symmetry (NCS) restraints were applied in Phenix to ensure accurate fitting and maintain structural integrity in these less-resolved regions. The model statistics were validated using MolProbity^60^. Structural figures were made using UCSF ChimeraX and PyMOL^61^ and UCSF Chimera^62^.

### Structural analyses

All structural analyses were performed using UCSF ChimeraX. Buried surface areas and inter-subunit molecular interactions were calculated using the PISA (Protein Interfaces, Surfaces and Assemblies) server (https://www.ebi.ac.uk/pdbe/pisa/). For buried surface area calculations, a model was generated in which the residue numbering of each protomer was corrected to begin at residue 1 (rather than 0, −1, −2, etc., as originally assigned during model building), allowing the buried surface area between the two tiers of the assembly to be measured accurately. All structural figures were generated using UCSF ChimeraX and PyMOL (Schrödinger, LLC). The radius of the central translocation tunnel was calculated using Caver Analyst 2.0.

### Multiple sequence alignment and phylogenetic analysis

Sequences of homologous structures were retrieved following a BLAST search. Multiple sequence alignment and phylogenetic tree construction were performed in Jalview^63^ using the MUSCLE alignment algorithm^64^.

### Isothermal Titration Calorimetry (ITC)

Isothermal titration calorimetry (ITC) experiments were performed using a Nano ITC microcalorimeter instrument (TA Instruments). Purified YsaN protein was dialyzed against buffer containing 30 mM HEPES (pH 7.5), 150 mM NaCl, and 5 mM MgCl₂ for 12 h with three buffer exchanges. All ligands were prepared in the same dialysis buffer to minimize buffer mismatch. Two independent ITC experiments were conducted to characterize nucleotide binding. For ATPγS titrations, YsaN was loaded into the sample cell at a concentration of 0.014 mM, and ATPγS was placed in the injection syringe at 0.67 mM. Titrations consisted of 30 sequential injections of 3.5 μL ligand. For ADP titrations, YsaN was used at 0.014 mM, with ADP at 0.85 mM in the syringe, and titrations consisted of 30 injections of 3.5 μL ligand. All measurements were performed at 283 K with continuous stirring of 125 RPM, with 300 s intervals between successive injections. Blank runs were performed by titrating ligands into buffer alone, under identical conditions, and were deducted from the protein-ligand titration prior to analysis. Binding isotherms were analyzed by nonlinear least-squares fitting using NanoAnalyze software (TA Instruments). The data were fit to a single-site binding model to obtain the binding stoichiometry (n), dissociation constant (Kd), and enthalpy change (ΔH). The Gibbs free energy change (ΔG) was calculated using ΔG = RT ln Kd, where R is the universal gas constant and T is the absolute temperature. The entropy change (ΔS) was derived from the relationship ΔG = ΔH - TΔS. Three independent titrations were performed for all experiments.

### Molecular Dynamics (MD) simulations System setup and preparation

The cryo-EM structures of the YsaN dodecamer (12-mer) and tetradecamer (14-mer) served as the initial coordinates for molecular dynamics (MD) simulations. To generate the nucleotide-bound states, ATP and ADP coordinates were mapped into the structures using ATP- and ADP-bound YsaN protomer models predicted by the AlphaFold Server. Specifically, each experimental YsaN protomer was structurally aligned to the AlphaFold models using UCSF ChimeraX to transfer the respective nucleotide coordinates. Because the experimental higher-order assemblies were resolved utilizing non-crystallographic symmetry (NCS) refinement (yielding essentially identical protomers), this nucleotide-bound monomer was systematically superimposed onto each protomer within the 12-mer and 14-mer cryo-EM maps. This protocol yielded six independent simulation systems: apo, ATP-bound, and ADP-bound 12-mer assemblies; and apo, ATP-bound, and ADP-bound 14-mer assemblies.

All structures were prepared utilizing the Protein Preparation Wizard within Schrödinger Maestro (Release 2025-4; Schrödinger, LLC, New York, NY, USA). Bond orders were reassigned, missing side-chain hydrogen atoms were added, and protonation states for ionizable residues and ligands were assigned at pH 7.4 ± 2.0 using Epik. Hydrogen-bonding networks were optimized via PROPKA. Subsequently, restrained energy minimization was performed using the OPLS4 force field until a maximum heavy-atom root-mean-square deviation (RMSD) of 0.30 Å was achieved.

### Simulation parameters

Simulation systems were constructed using the Desmond^65^ System Builder module (Schrödinger Release 2025-4). Each protein complex was solvated in an orthorhombic periodic simulation box containing explicit TIP3P^66^ water molecules. The systems were neutralized with the addition of counterions, and NaCl was supplemented to achieve a physiological ionic strength of 0.15 M.

All MD simulations were executed using the Desmond MD engine with the OPLS4^67^ force field. Following the default Desmond relaxation protocol, 200 ns production simulations were carried out under the isothermal–isobaric (NPT) ensemble^68^ at 300 K and 1.01 bar. The system temperature was regulated using a Nosé–Hoover chain thermostat^69^, and pressure was maintained using a Martyna–Tobias–Klein barostat^70^. Long-range electrostatic interactions were computed utilizing the particle mesh Ewald (PME) method^71^, while short-range van der Waals and electrostatic interactions were truncated at a 9 Å cutoff distance. Periodic boundary conditions were applied in all three dimensions. The equations of motion were integrated using a 2 fs time step, with trajectory coordinates recorded every 200 ps for subsequent analyses.

### Trajectory analysis

Simulation trajectories were primarily analyzed using the Desmond Simulation Interaction Diagram module. Global structural stability was evaluated by calculating the backbone RMSD and residue-wise root-mean-square fluctuation (RMSF). Time-dependent secondary structure evolution was tracked using the built-in secondary structure analysis module. Protein–nucleotide interaction profiles—including hydrogen bonds, hydrophobic interactions, ionic contacts, and water-mediated bridges—were quantified to determine interaction occupancies throughout the simulation. The radius of gyration ( ) and dynamic cross-correlation matrices (DCCM) were computed using custom Python scripts.

### Principal Component Analysis (PCA) and Free Energy Landscape (FEL)

To capture the dominant conformational motions of the 12-mer and 14-mer complexes, PCA was performed on the C- atoms using the scikit-learn library in Python. To ensure proper structural equilibration, the initial 50 ns (250 frames) of each trajectory were discarded. The remaining frames were translationally and rotationally aligned to their respective first equilibrated frame via Kabsch superposition. The covariance matrix of atomic fluctuations was diagonalized to yield the principal components (PCs), and the first two components (PC1 and PC2)—representing the largest variance in structural dynamics— were retained for dimensionality reduction.

The Free Energy Landscape (FEL) was constructed by projecting the aligned trajectories onto the 2D subspace defined by PC1 and PC2. A 2D probability density histogram (100 × 100 bins) was generated from these projections and subsequently smoothed using a Gaussian filter (σ = 1.5 bins) to yield a continuous landscape. Probabilities were converted into Gibbs free energy (Δ*G*) using the Boltzmann relationship:

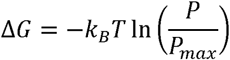

Where *k_B_* is the Boltzmann constant, is the simulation temperature (300 K), *p* is the probability of a specific conformational state, and *p_max_* is the probability of the most highly populated state. The lowest energy basins on the resulting plots correspond to the most stable and frequently sampled structural conformations. All MD simulation plots were generated using Python scripts.

## Author Contributions

B.B., B.C., S.D. and D.P. wrote and edited the paper. Cryo-EM grid vitrification, data collection, and data processing were done by B.B. & D.P. Model building and structural analysis were performed by B.B. & B.C. Protein purification and Biochemical studies were carried out by B.C., B.B and S.N. MD Simulations and analysis were carried out by B.B. & D.P. The project was supervised by S.D. & D.P. All authors approved the final copy of the paper.

## Classification

**Major Category:** Biological Sciences, **Minor Category 1:** Microbiology, **Minor Category 2:** Biophysics and Computational Biology

## Data Availability

EM maps of the YsaN 11mer, 12mer, 13mer, and 14mer filament assemblies were deposited to the EM Data Bank under accession codes EMD-65882, EMD-66223, EMD-65920 and EMD-66331, respectively. The corresponding atomic models were deposited to the PDB under accession codes 9WD4, 9WTO, 9WF0 and 9WX6, respectively. The individual focused maps for the monomer of 12mer and 14mer were deposited to the EM Data Bank under accession codes EMD-66220, and EMD-66329, respectively.

## Acknowledgements

Cryo-EM data were collected at the CSIR-CCMB EM Facility. Computational resources and support were provided by the high-performance computing facilities at CSIR-4PI. The authors also acknowledge the Central Instrumentation Facility of IICB for providing access to Central instruments and Schrödinger software used in this study.

