## Supporting Information for "Cryo-EM structures of the YsaN ATPase reveal dodecameric and tetradecameric assemblies in the *Yersinia enterocolitica* T3SS"

**This PDF file includes:**

Figures S1 to S7
Legends for Movies S1 to S8

**Other supporting materials for this manuscript include the following:**

Movies S1 to S8

Figures

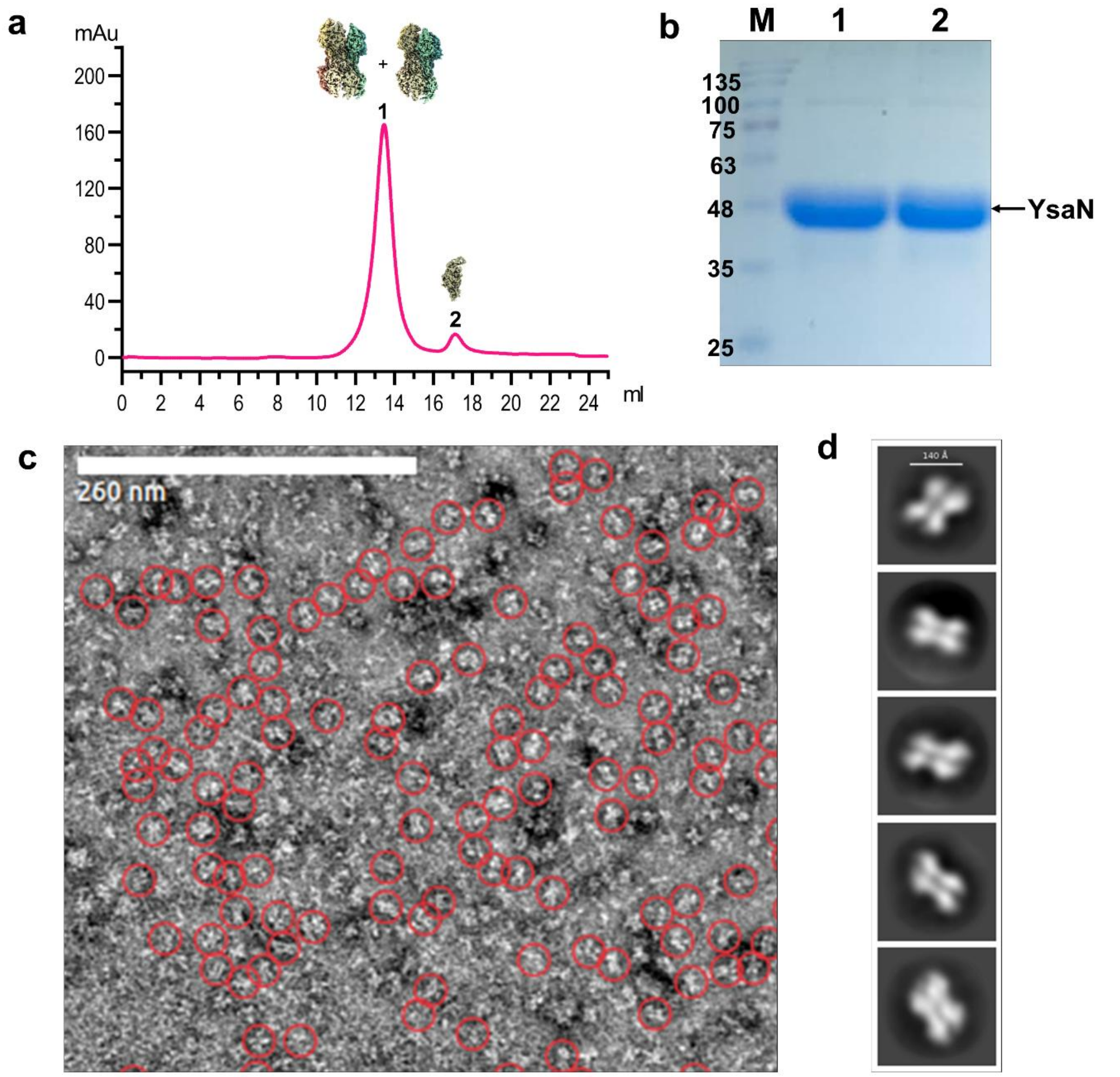


**Fig. S1. Purification and negative-stain electron microscopy of YsaN. a**, Size-exclusion chromatogram of the YsaN complex utilizing a Superose 6 Increase 10/300 GL column, show two different peaks, one eluting around 13.4 ml, corresponding to higher order oligomer (12mer and 14mer) and the other at 17.06 mL corresponding to monomer. **b**, SDS-PAGE analysis of the corresponding peaks. **c**, Representative negative-stain micrograph highlighting well-dispersed YsaN particles (red circles). **d**, Reference-free 2D class averages.

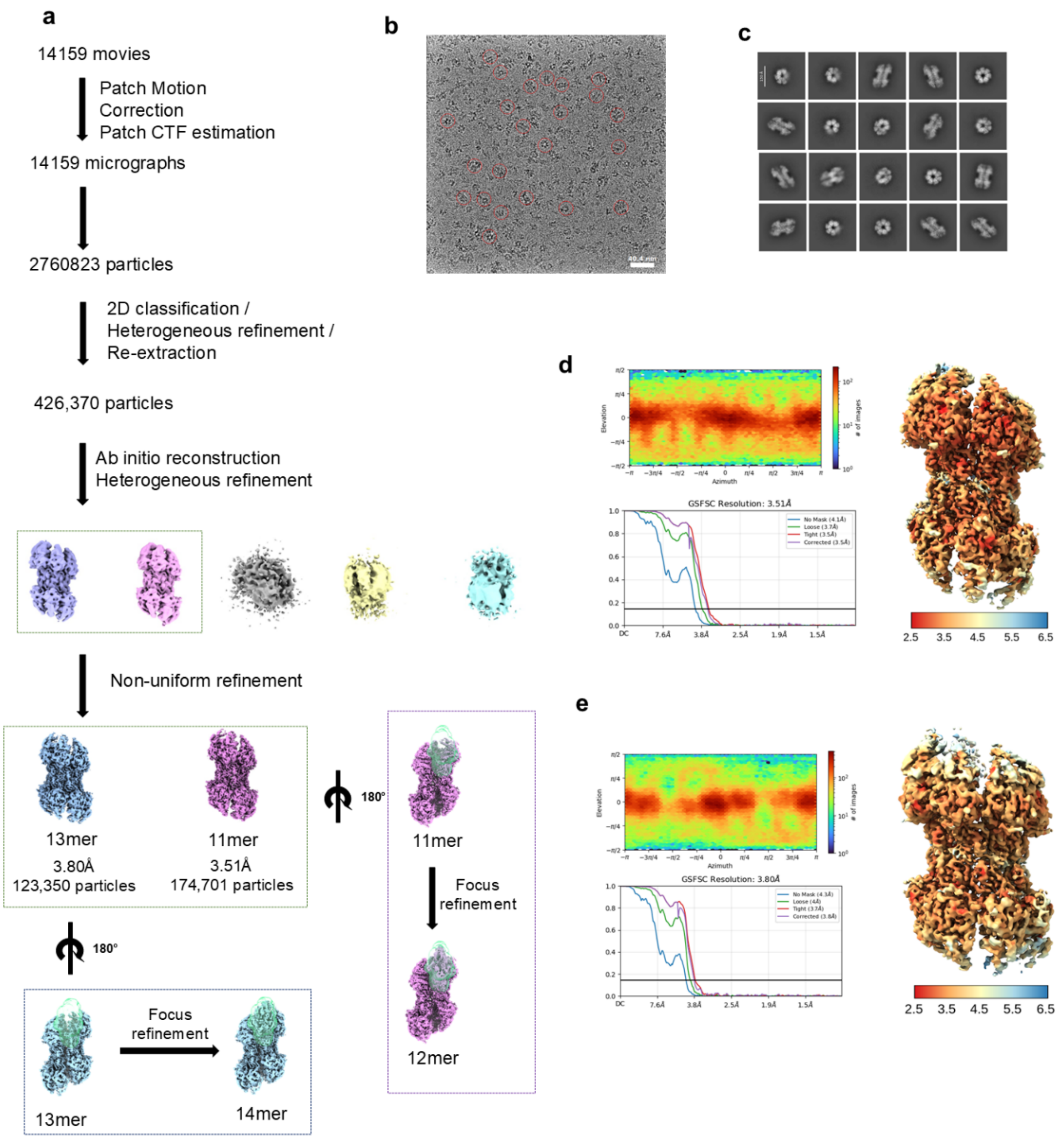


**Fig. S2. Overall data processing workflow used for 3D reconstruction of the complexes. (a)** Overall Cryo-EM data processing overflow, **(b)** Representative micrograph, **(c)** Selected reference-free 2D class averages. **(d)** Orientation distribution, GS-FSC curve and local resolution estimation of YsaN dodecamer and **(e)** tetradecamer.

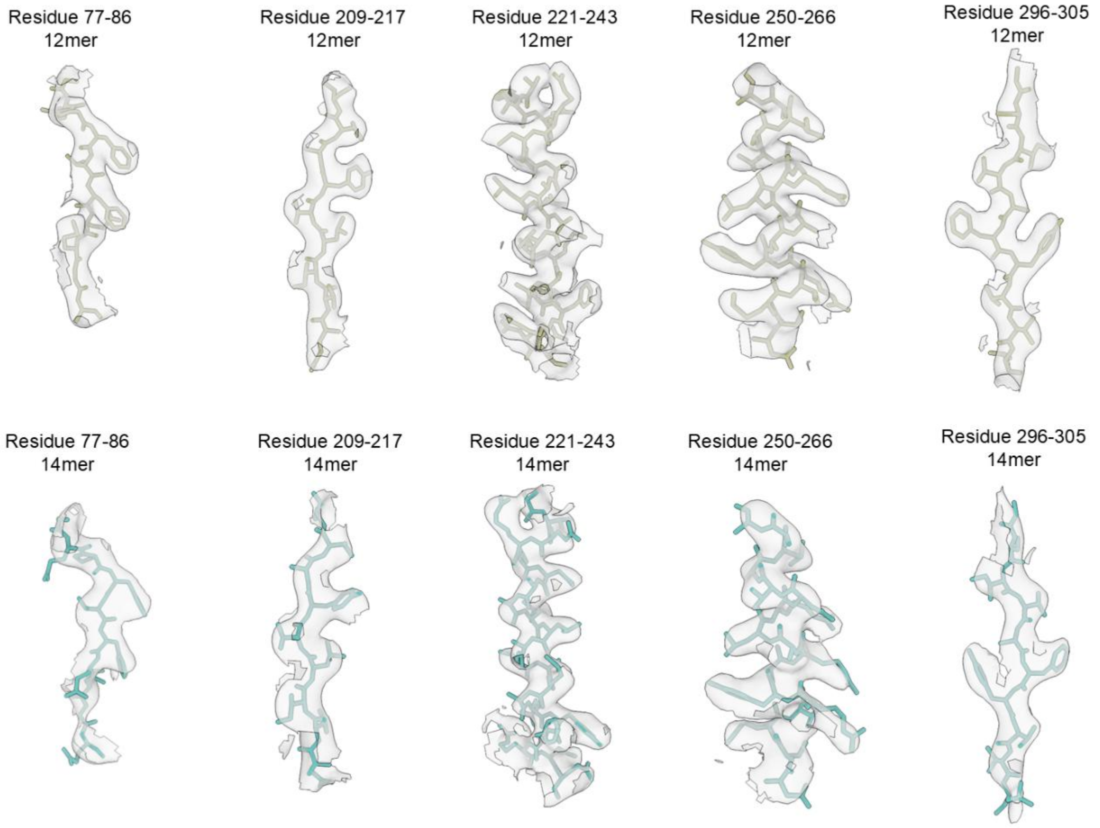


**Fig. S3. Representative cryo-EM density maps of the YsaN complex.** The densities of YsaN are contoured at 6.70 σ.

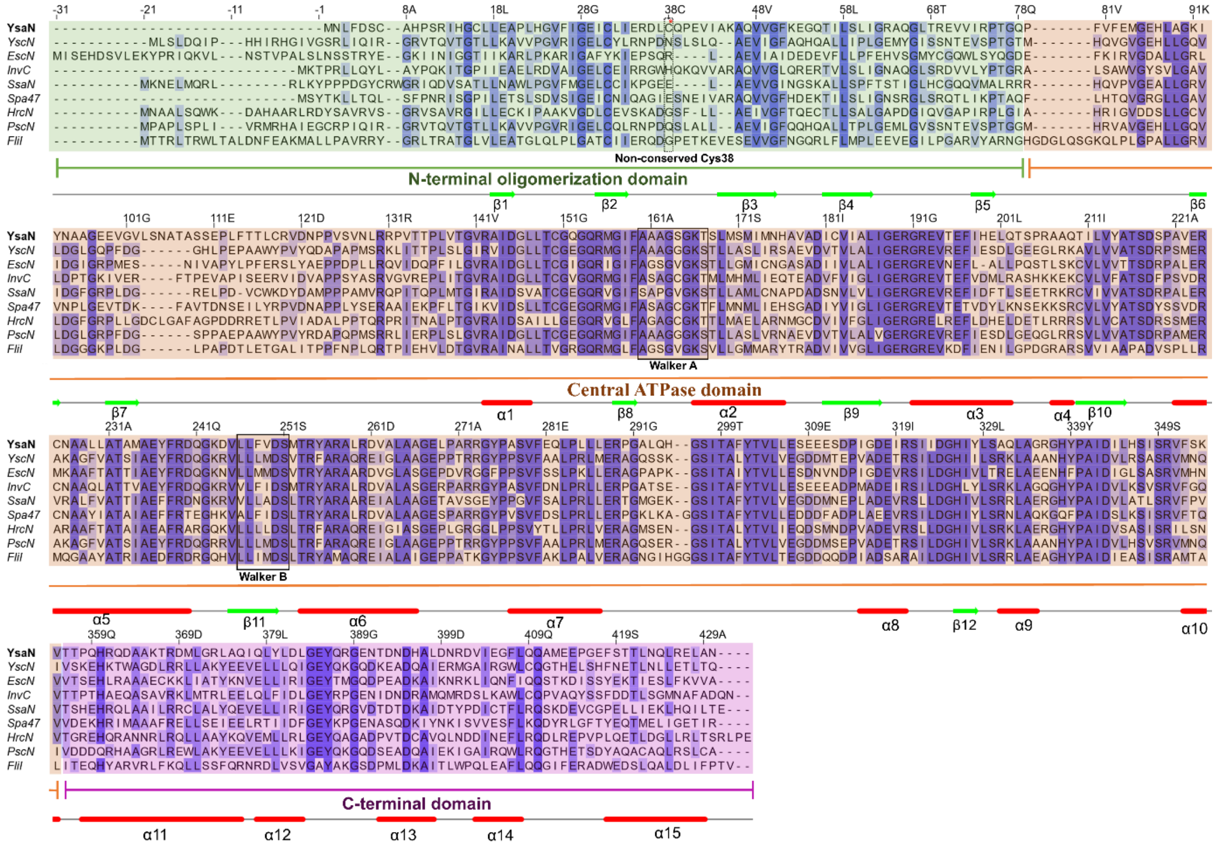


**Fig. S4. Multiple sequence alignment of YsaN with its homologs ATPases.** Multiple sequence alignment of YsaN with homologous ATPases, including YscN (*Yersinia pestis*; Q7ARI8), EscN (*Escherichia coli* EPEC; B7UMA6), InvC (*Salmonella enterica* serovar Typhimurium; P0A1B9), SsaN (*Salmonella enterica*; P74857), Spa47 (*Shigella flexneri*; P0A1C1), HrcN (*Pseudomonas syringae*; Q8RK01), PscN (*Pseudomonas aeruginosa*; A0A0H2Z9F1), and FliI (*Salmonella enterica*; P26465). The alignment was generated using MUSCLE and visualized in Jalview. Secondary structure elements derived from the YsaN structure are displayed below the alignment. Conservation is indicated by a blue color gradient, where deep blue represents highly conserved residues, and white indicates no conservation. The three structural domains of YsaN are indicated by colored bars above the secondary structure: the N-terminal domain (NTD; green), the central ATPase domain (orange), and the C-terminal domain (CTD; pink). The Walker A (P-loop) and Walker B motifs are highlighted, showing strong conservation across all homologs. In contrast, the cysteine residue implicated in disulfide bond formation in YsaN is not conserved among orthologs.

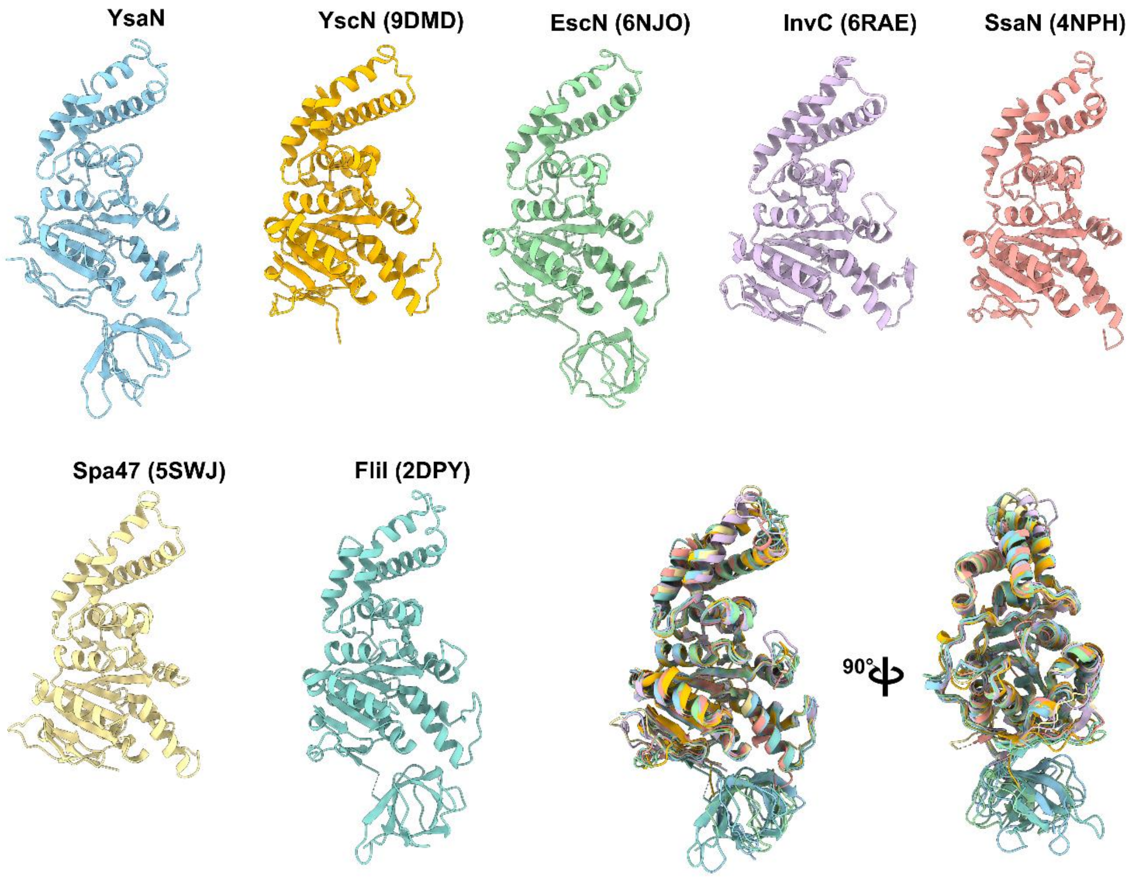


**Fig. S5. Structural superposition of the full-length YsaN protomer with homologous T3SS and flagellar ATPases.** Cartoon representations of the YsaN protomer (light blue) alongside the available experimental structures of related homolog ATPases: YscN (PDB: 9DMD, orange), EscN (PDB: 6NJO, green), InvC (PDB: 6RAE, purple), SsaN (PDB: 4NPH, salmon), Spa47 (PDB: 5SWJ, pale yellow), and the flagellar ATPase FliI (PDB: 2DPY, teal). The bottom right panels display a global structural alignment of all seven protomers, presented in two orthogonal views related by a 90° rotation. The structural overlay demonstrates the strict evolutionary conservation of the central AAA+ catalytic core and the C-terminal domains. Crucially, the superposition visually highlights the presence of the intact N-terminal oligomerization domain in the full-length YsaN model, a critical structural element that is truncated or entirely absent in several of the previously reported homologous structures.

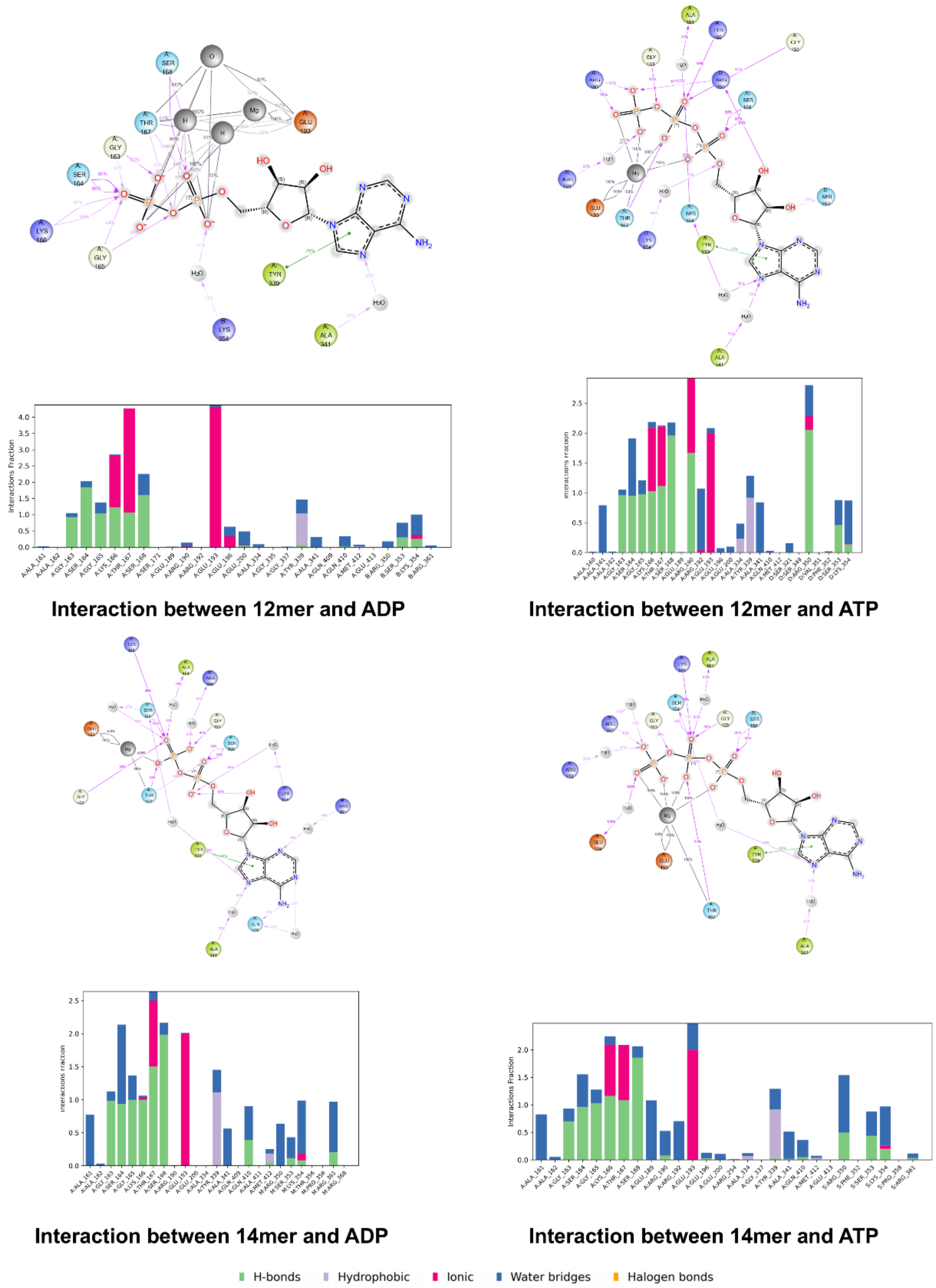


**Fig. S6. Nucleotide interaction profiling of YsaN assemblies.** Top and Middle: Detailed 2D ligand interaction diagrams and corresponding protein-ligand interaction fraction histograms for the 12-mer and 14-mer YsaN complexes bound to ADP and ATP, extracted from the molecular dynamics simulations. The 2D diagrams map the specific atomic contacts between the nucleotide and the binding pocket. The histograms quantify the fractional occupancy of these interactions (e.g., hydrogen bonds, hydrophobic contacts, ionic interactions, and water bridges) over the simulation trajectory, highlighting the dominant coordination by the conserved Walker A motif (residues 160–167) and the stabilizing stacking interaction provided by Tyr339.

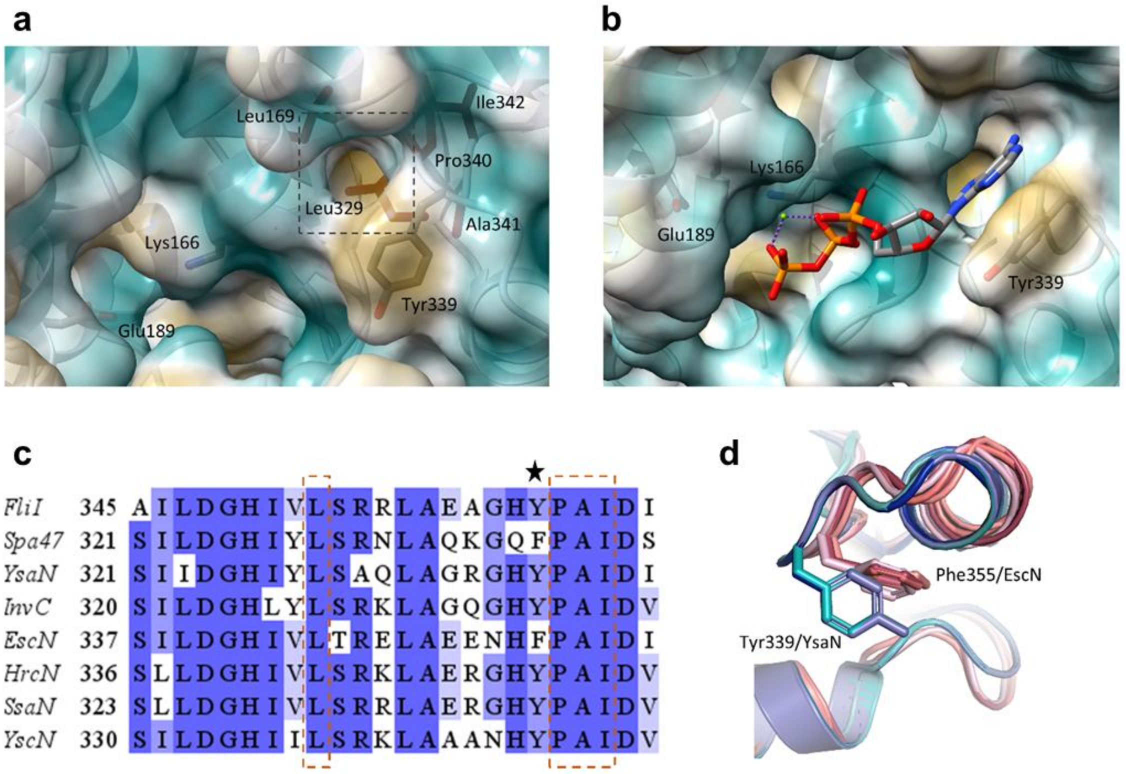


**Fig. S7. Conserved Tyr339 occludes the adenine binding hydrophobic pocket in YsaN.**
**a)** The closed adenine binding hydrophobic pocket in YsaN 12-mer structure (PDB ID: 9WD4). The surface is coloured by higher (yellow) to lower (blue) hydrophobicity. The position of catalytic glutamate (Glu189) and P-loop lysine (Lys166) is shown by stick representation. The conserved hydrophobic residues lining the pocket are shown by stick representation. **b)** Hydrophobic surface representation of ATP-docked and energy minimized YsaN 12-mer structure showing opening of the adenine binding pocket by movement of the Tyr339 sidechain upon ligand binding. **c)** Multiple sequence alignment of T3SS ATPases across species shows conservation of the hydrophobic pocket residues, the Leu329, Pro340, Ala341, and Ile342 lining the pocket are highlighted by orange dashed box. The conserved aromatic residue responsible for stacking the adenine group is marked by a black star. **d)** Superposition of EscN and YsaN subunits show rotation of the aromatic side chain of Tyr339 compared to Phe355 of EscN. The EscN subunits are coloured in shades of pink, YsaN subunits are coloured in shades of blue.

**Movie S1 (separate file). Structures of the dodecameric (12mer) and tetradecameric (14mer) YsaN assemblies.** A comparative 360-degree rotation of the full-length YsaN 12-mer (left) and 14-mer (right) complexes. The video initially displays the global cryo-EM density maps and subsequently shows the atomic models, with individual protomers distinctly coloured.

**Movie S2 (separate file). The helical rise in the YsaN assembly.** This animation sequentially traces the continuous helical progression of the 12-mer YsaN complex. The N-terminal domain of each protomer is distinctly coloured and shown one after another to clearly highlight the steady vertical translation and rotational twist that define the helical-like architecture. In the final segment of the video, the complete structure of the complex is shown, with the core ATPase and C-terminal domains rendered as transparent white outlines.

**Movie S3 (separate file). Molecular dynamics simulations of the 12mer YsaN assembly.** The structural trajectories of the YsaN 12-mer complex across three distinct nucleotide states: apo (left, blue), ADP-bound (centre, orange), and ATP-bound (right, green). The respective nucleotide ligands are visible as distinct space-filling models bound within the active sites of the ADP-bound and ATP-bound assemblies.

**Movie S4 (separate file). Molecular dynamics simulations of the 14mer YsaN assembly.** Similar structural trajectories of the YsaN 14-mer complex across three distinct nucleotide states: apo (left, blue), ADP-bound (centre, orange), and ATP-bound (right, green). The respective nucleotide ligands are visible as distinct space-filling models bound within the active sites of the ADP-bound and ATP-bound assemblies.

**Movie S5 (separate file). Dominant principal motion (PC1) of the 12mer YsaN assembly.** The first principal component (PC1) extracted from the molecular dynamics trajectories of the 12mer YsaN complex. Displayed simultaneously in side, top, and bottom views.

**Movie S6 (separate file). Secondary principal motion (PC2) of the 12mer YsaN assembly.** The second principal component (PC2) extracted from the molecular dynamics trajectories of the 12mer YsaN complex. Displayed simultaneously in side, top, and bottom views.

**Movie S7 (separate file). Dominant principal motion (PC1) of the 14mer YsaN assembly.** The first principal component (PC1) extracted from the molecular dynamics trajectories of the 14mer YsaN complex. Displayed simultaneously in side, top, and bottom views.

**Movie S8 (separate file). Secondary principal motion (PC2) of the 14mer YsaN assembly.** The second principal component (PC2) extracted from the molecular dynamics trajectories of the 14mer YsaN complex. Displayed simultaneously in side, top, and bottom views.
